# Reward-based prosocial choices in mice

**DOI:** 10.1101/2025.02.21.639526

**Authors:** Joan Esteve-Agraz, Víctor Javier Rodríguez-Milan, Cristina Márquez

## Abstract

Prosocial behaviors, actions that benefit others, are an essential part of the social life of humans and other animals, by promoting bonding and cohesion among individuals and groups. Here we present a new behavioural paradigm to assess prosociality in food foraging contexts in laboratory mice, based in our paradigm previously developed for rats. In this task, the decision-maker can choose between two options, one that will only provide rewards to itself (selfish choice) or one that will reward both itself and its cagemate (prosocial choice). Our work reveals that prosocial choices in male mice are not widespread and are only observed in a small proportion of animals. Using detailed analysis of behavior, we describe that recipients of help express different social cues in prosocial and selfish trials, but decision makers do not take them into account to guide their choices. Furthermore, we assess how the level of individual training and the physical layout of the paradigm might affect the performance in this social task. Only those mice with increased social attention (16% of the animals) display prosocial preferences, suggesting these to be rooted in similar behavioural factors and social interactions that we previously described in other work with rats.

## Introduction

Prosocial behaviors, defined as actions that benefit others, constitute a fundamental aspect of the social dynamics supporting positive social interactions and cohesion among individuals (Penner. et al., 2005; Gachomba, Esteve-Agraz et al., 2024). The study of prosociality extends beyond humans as these actions are conserved across different species, offering valuable insights into the evolutionary roots and underlying mechanisms that govern such social interactions (Cronin, 2012; Rault, 2019; Takimoto-Inose, 2021). Over the last years, a growing number of experimental works has started to recognize the responsiveness of rodents to the emotional displays of conspecifics (for review see Hernandez-Lallement et al., 2022; Keysers et al., 2022; Pérez-Manrique & Gomila, 2022; Gachomba, Esteve-Agraz, et al., 2024). Most of these findings are related to prosocial actions towards distressed conspecifics (i.e. in pain, fear or stress), which are very robust emotional responses in rodents (Burkett et al., 2016; C.-L. Li et al., 2018; L.-F. Li et al., 2019; Wu et al., 2021; Bartal et al., 2011; Sato et al., 2015; Breton et al., 2022; Hernandez-Lallement et al., 2020; Song et al., 2023; see Keysers & Gazzola, 2023, for review). The study of prosocial behaviours within positive contexts (i.e. non-distressful) has received comparatively less attention (Gachomba, Esteve-Agraz, et al., 2024), hindering a more comprehensive understanding of the multifaceted nature of social cognition and prosocial actions within these species. Yet, there have been important contributions showing that rodents are also prosocial in reward-based contexts, as demonstrated, for instance, with the prosocial choice task (PCT). This paradigm has been established to measure affiliative behaviors such as other-regarding preferences for reward provision in different taxa (Burkart et al., 2007; De Waal et al., 2008; Cronin et al., 2009; Massen et al., 2010; Schwab et al., 2012; Péron et al., 2013; Quervel-Chaumette et al., 2015; Márquez et al., 2015; Gachomba et al., 2022). Studies using different versions of this paradigm have shown that rats are prosocial, in most conditions, by preferring those choices that reward a conspecific (Márquez et al., 2015; Hernandez-Lallement et al., 2015; Kentrop et al., 2020; Joushi et al., 2022; Gachomba et al., 2022). Evidences for reward-based prosocial actions in mice are still scarce, and results somehow contradictory (Scheggia et al., 2022; Misiołek et al., 2023), making it hard to find common behavioural and neural mechanisms delimiting the extent of prosocial choices in these two rodent species.

Testing with different paradigms to address similar questions might result in conflicting evidence but is also highly relevant and beneficial for the advancement of our knowledge. Using the same paradigm to compare across different species would further provide valuable insights. Therefore, the development and application of paradigms that allow for an objective and standardized dissection of the key details that are intrinsic to the complexity of social behaviours, and the comparison across species, will greatly benefit the research into the mechanisms of prosociality. In the present work, which aims to provide a framework for studying the proximal behavioural factors underlying prosocial actions, we introduce a new behavioural paradigm to assess prosocial decision-making in mice. This paradigm is based on a previous paradigm developed for rats, in which our group showed that Sprague-Dawley rats behave prosocially towards conspecifics in a prosocial choice task within a food-foraging context (Márquez et al., 2015; Gachomba et al., 2022). Briefly, the task consists in a double T-maze, in which a decision-maker (*focal* animal) controls the access to food rewards for itself and the recipient animal. In each trial, the focal animal can either choose the prosocial option (i.e., by providing access to food for itself and the recipient) or the selfish option that provides food access only to itself. Recipient animals in this task are not just passive agents; they actively seek to access their rewarded arm, and these displays of food-seeking behavior are necessary for the emergence of prosocial choices by focal rats (Márquez et al., 2015).

Here we assessed whether C57BL/6J laboratory male mice showed the affective/cognitive capacities to display prosocial choices, by providing food to their conspecifics, as observed in rats, using a behavioral task adapted from the original one developed for rats. We found that most mice displayed no clear preferences to provide food to conspecifics, although marked individual differences were observed. Our results suggest that although mice do not have a tendency for widespread prosociality, there are some individual factors and social interactions that promote the emerge of prosocial biases, which interestingly, are the same ones previously described to be necessary for prosociality in rats. This research, aimed to expand and complement our scarce knowledge of reward-based prosociality in mice, provided a unique opportunity to elucidate species-specific differences in social cognition and behaviour, and to further pave the way for developing comparative studies of the neural correlates of prosociality.

## Results

### Study of reward-based prosocial choices in mice

#### Prosocial choices in reward-based T-maze task are not widespread in C57BL/6 male mice

In previous work with rats, our group demonstrated that (1) decision-maker rats are sensitive to the food-seeking behavior displayed by the recipient animals prior to choice (Márquez et al., 2015). This is necessary for the emergence of a prosocial preference, but not sufficient, as (2) information about the reward contingencies of the recipient was also relevant for prosocial choices to emerge. Keeping these two important mechanisms in mind, we developed a fully automated double T-maze (Figure 1a), which decreased the possible interferences by the experimenter and provided a precise and controlled monitoring of the behaviour of the interacting mice (Supplementary Video 1). The configuration of this maze separated spatially and temporally the moments of decision from those of reward delivery (Figure 1a-c). Each of the individual T-mazes (one per animal) contained a central ‘choice area’, where two nose ports for each animal were located, used for displaying food-seeking behaviour and decisions (i.e. by nose-poking into the IR ports). The central zone gave access to two lateral areas, gated by automatic doors, where mice retrieved the rewards from a food receptacle according to the contingencies of the task. After reward retrieval, the doors opened allowing the animals to go back to the choice area to start a new trial. The two individual T-mazes were connected by a perforated and transparent partition, which allowed mice to exchange different sensory information in the choice area as well as in the reward areas. To ensure that focals had the opportunity to perceive the displays of food-seeking behaviour and the reward retrieval by the recipient, the nose-ports of the focal mouse were active only after the recipient poked at least once in any of the IR ports (i.e. indicating the recipients interest and presence near the choice ports) and the food pellet of the recipient was delivered only when the focal mouse was present in the reward area.

**Figure 1.**
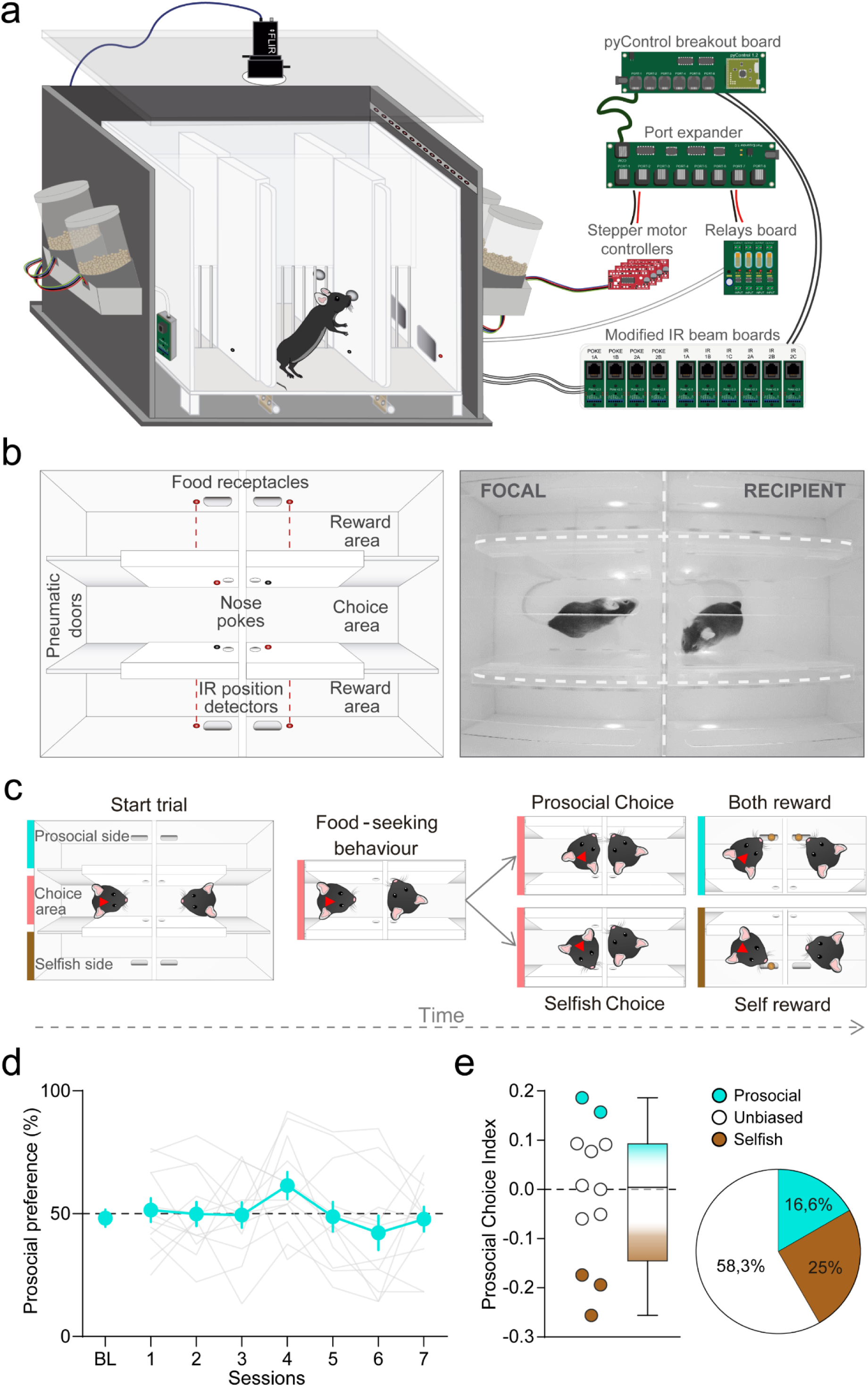
Mice prosocial choices in double T-maze. **a**. Hardware and peripherals used for the assessment of prosocial decision-making with a double-T maze arena. The arena is located inside a sound attenuation box and illuminated with IR and dim white light to enable high quality video recordings. The setup is made of laser-cut white acrylic treated to avoid reflections from the IR camera placed on the box centre-top. Custom-made pellet-dispensers hold outside the sound attenuation box to reduce head and noise. Pneumatic cylinders are below the base of the arena providing smooth gating of the doors. **b**. Hardware position schematic (left) and real top image (right) of the maze-based setup configuration. The T-mazes are joined by a perforated and transparent partition. For each side there is a central choice area with two nose-ports located in each wall (for decisions and displays of food-seeking behaviour), and modified IRs to detect the mice position. There are acrylic doors connected to pneumatic cylinders at the end of the corridor that give access to the reward areas. In these zones food pellets are delivered by automated food-dispensers located outside the arena. **c**. Timeline structure of prosocial choice task. Trials start with both animals in the choice area, the recipient will display food-seeking behaviour by nose-poking into any of the ports which will activate the decision ports of the focal mouse (red triangle in the head). The focal then will decide to go either side of the maze by nose-poking in any of the ports. Poking into the prosocial port will deliver a food-pellet to both animals while choosing the selfish port will only deliver a pellet for the focal and none to the recipient. The different separated areas are colour-coded (choice area: pink, prosocial side: blue, selfish side: brown). **d**. Prosocial preference of mice in maze-based arena over the seven testing sessions. BL refers to baseline, used to evaluate individuals’ preference in the last two sessions of individual training. Blue thick line corresponds to mean±SEM, grey lines correspond to each individual. At population level, animals did not display any preference for prosocial or selfish choices. **e**. Distribution of Prosocial Choice Indexes to study individual differences in prosociality. Positive values show a preference for the prosocial option, negative values indicate preference for the selfish option, and values close to 0 indicate chance preference. Blue dots correspond to prosocial mice, grey dots are unbiased and brown selfish. On the right, pie chart: distribution of mice after permutation test of Prosocial Choice Indexes

We tested pairs of mice in our PCT, where a decision-maker mouse (focal) could choose to provide food reward only to itself (*selfish option*) or to itself and the recipient mouse (*prosocial option*) (Figure 1c). Before social testing, mice were individually trained for instrumental learning and maze navigation (Supplementary Figure 1). By the end of the individual training, decision makers showed no general side bias (one sample t-test against chance (50), t_(11)_=-0.489, p=0.634, BF_10_=0.319) (Supplementary Figure 1c) and recipient animals showed clear displays of food seeking behavior (poke specificity toward the prosocial side: 94.11% ± 1.01) (Supplementary Figure 1f). Then, focal and recipient mice were tested together in the PCT, where reward delivery for the two animals depended on the focals’ choices. For this set of experiments, 12 pairs of male mice (C57BL/6) underwent 7 sessions of 30 minutes of the PCT. Mice were not food-restricted during prosocial testing days, to avoid possible stress-related influence on prosocial tendencies.

During the social task, decision-makers did not develop a preference for prosocial or selfish options over sessions (repeated measures ANOVA with ‘session’ as within subjects: F_(6)_ = 1.857, p=0.102, BF_incl_= 0.637) (Figure **Error! Reference source not found.**1d). These results suggest that mice had no preference for choosing the option that delivered food to their conspecifics, against what we had observed in rats. However, most of the animals changed substantially their preferred choices over sessions (increased and decreased), showing high individual variability. In order to account for the differences in prosociality between individuals, we computed a Prosocial Choice Index (PCI) (see Statistical Analysis in Methods). Positive PCIs reflect a change towards a prosocial preference from chance (PCI=0). We found moderate evidence supporting the lack of preference for any of the choices (one sample t-test: t_(11)_=-0.251, p=0.59, BF_+0_=0.242). A permutation test on the PCI of the individuals revealed that out of 12 mice, 2 were considered prosocial, 7 unbiased and 3 selfish (Figure 1e, Table 1).

**Table 1.** Chance interval bounds generated by permutation test for each pair. Related to Figure 1, 7 and 8. Low and high bounds show the 95% confidence interval for each focal mouse.

| Standard PCT in double T-Maze |  |  |  |  |  |  |
| --- | --- | --- | --- | --- | --- | --- |
| Pair # | 1 | 2 | 3 | 4 | 5 | 6 |
| Lower bound | -0,096 | -0,164 | -0,179 | -0,152 | -0,128 | -0,152 |
| Upper bound | 0,096 | 0,194 | 0,179 | 0,131 | 0,128 | 0,152 |
| Pair # | 7 | 8 | 9 | 10 | 11 | 12 |
| Lower bound | -0,115 | -0,131 | -0,136 | -0,115 | -0,140 | -0,140 |
| Upper bound | 0,115 | 0,131 | 0,136 | 0,115 | 0,140 | 0,140 |
| Low training PCT in double T-Maze |  |  |  |  |  |  |
| Pair # | 1 | 2 | 3 | 4 | 5 | 6 |
| Lower bound | -0,200 | -0,175 | -0,192 | -0,176 | -0,165 | -0,186 |
| Upper bound | 0,200 | 0,175 | 0,192 | 0,176 | 0,165 | 0,186 |
| Pair # | 7 | 8 | 9 | 10 | 11 | 12 |
| Lower bound | -0,263 | -0,273 | -0,278 | -0,185 | -0,176 | -0,257 |
| Upper bound | 0,263 | 0,273 | 0,278 | 0,185 | 0,176 | 0,257 |
| Standard PCT in double chamber |  |  |  |  |  |  |
| Pair # | 1 | 2 | 3 | 4 | 5 | 6 |
| Lower bound | -0,063 | -0,067 | -0,099 | -0,067 | -0,078 | -0,099 |
| Upper bound | 0,063 | 0,072 | 0,099 | 0,067 | 0,078 | 0,099 |
| Bottom preference in double chamber |  |  |  |  |  |  |
| Pair # | 1 | 2 | 3 | 4 | 5 | 6 |
| Lower bound | -0,063 | -0,065 | -0,063 | -0,067 | -0,078 | -0,081 |
| Upper bound | 0,063 | 0,065 | 0,063 | 0,067 | 0,078 | 0,081 |

To improve our understanding of why focal mice did not prefer to choose the prosocial option, we analysed the behaviour of the animals according to their role during the social task. For this purpose, we performed a fine-grained analysis on the tracking data obtained by the animal pose estimation software DeepLabCut (DLC) and behavioural events extracted from pyControl (the software and hardware platform used to control these experiments).

#### Recipient mice display clear food-seeking behavior and reward related cues during the prosocial choice task

We first focused our analysis on understanding if the behaviour of recipient mice could explain the lack of choice preference found in focal mice. It has already been shown that the displays of food-seeking behaviour performed by the recipient rats are necessary for the emergence of a preference for the prosocial choices; therefore, we assessed if recipient mice performed clear food-seeking cues in the choice area (Figure 2a). To this end, we first quantified the number of pokes recipient mice did in each nose-port (prosocial and selfish ports) per trial (Figure 2b), and we found that the frequency of pokes into the prosocial port was much higher than those in the selfish nose-port (paired samples t-test: t_(11)_=7.723, p=9.120e^-6^, BF_10_=2341.38). We next calculated the poke specificity towards the prosocial port in both prosocial and selfish trials (Figure 2c), and observed that recipient mice poked almost exclusively towards the prosocial port independently on whether the focal would decide to be prosocial or selfish (paired samples t-test: t_(11)_=0.654, p=0.527, BF_10_=0.345). With tracking data obtained with DLC, we performed a ROI analysis and measured the amount of time that the head of recipient mice was detected inside the ROI around each of the nose-ports, as a less stringent measure of poke exploration (Figure 2d). We found that recipients spent a significantly higher amount of time near the nose port that gave access to the reward arms in comparison to the ‘selfish’ port (paired samples t-test: t_(11)_=4.654, p=7.005e-4, BF_10_=54.187).

**Figure 2.**
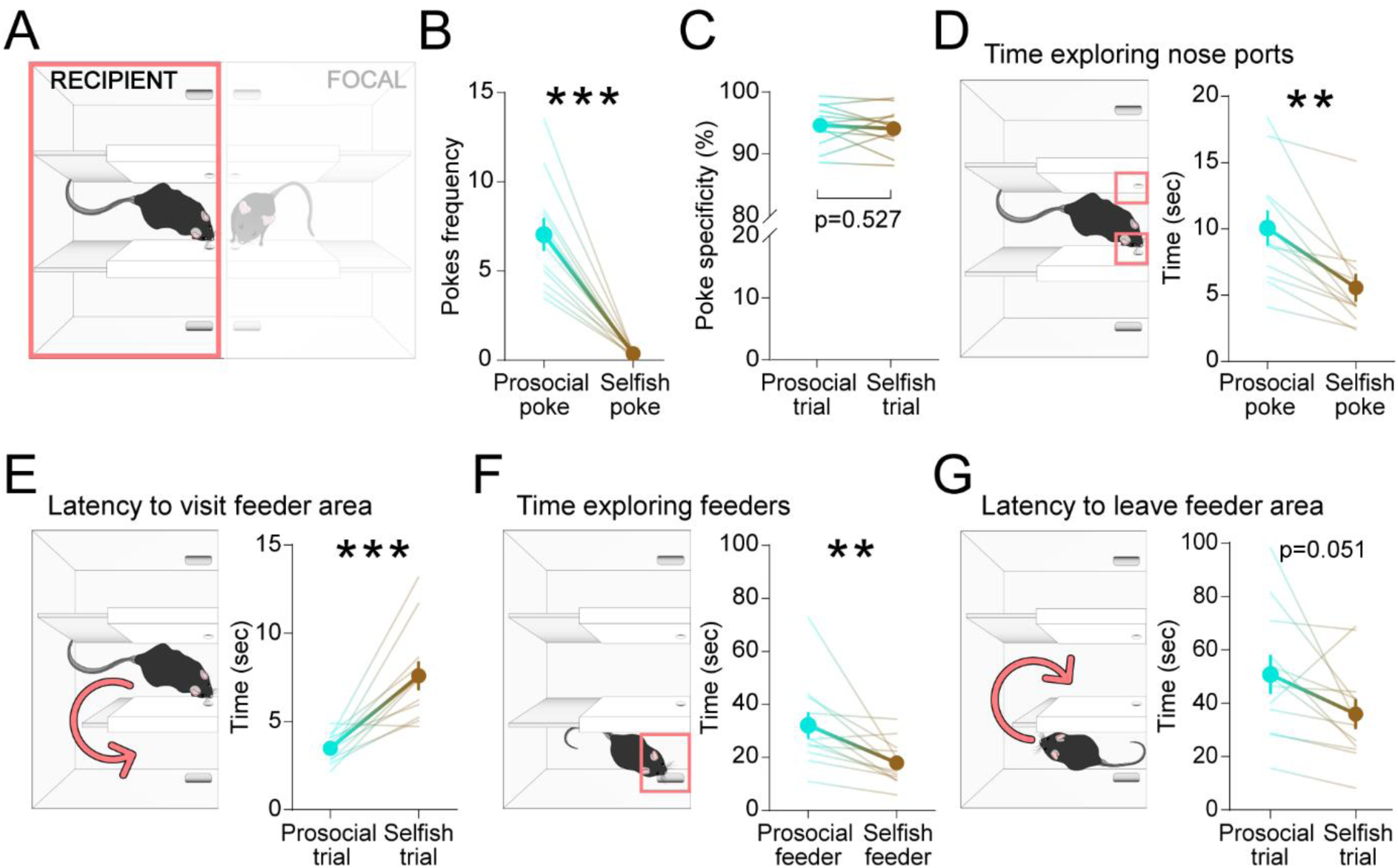
Recipient mice display food-seeking behaviour and react to reward contingencies. **a**. Illustration of the arena used for the PCT. The pink rectangle indicates that the following results are focused on recipients’ behavioural data. **b**. Nose-pokes frequency per trial. Quantification of pokes per trial done in the prosocial and selfish ports. Recipients do a significantly higher number of pokes in the prosocial nose-port compared to the selfish. **c**. Nose-poke specificity. For each recipient mouse, we calculated the proportion (in percentage) of pokes towards the prosocial port, both in prosocial trials (blue dots) and selfish trials (brown dots). Specificity is similar for prosocial and selfish trials, being around the 95% of pokes towards the prosocial port. **d**. Time exploring nose-ports. With recipients pose data, we performed a ROI analysis for the nose-ports (pink squares). We measured the nose label spent inside each of the two ROIs and found out that recipients spend almost double of the time exploring the prosocial port compared to the selfish**. e**. Latency to visit feeder areas. Time in seconds from choice to detection of the recipient mouse in the reward areas was significantly different. Recipient mice enter faster in the area where they get rewarded. **f**. Time spent exploring the feeders. Same as **d** for the area around the food receptacles (pink square). We also considered the detection of the head label to avoid data loss by occlusions from the walls separating the different areas. Recipients stayed significantly longer around the feeder where they are rewarded compared to the selfish. **g**. Latency to leave the reward areas. Ten seconds after reward delivery, automated doors opened to allow going back into the choice area to start a new trial. We found a tendency for recipients to leave the selfish reward area faster than the prosocial. For all graphs: degraded thicker line shows mean±SEM, thinner lines represent data from each individual. Blue = prosocial, brown = selfish.

Taken together, these results show that recipient mice actually displayed clear attempts to reach the food and access the rewarded arm. We then extended our analysis to the moments of the testing sessions that occur after the decision is made (and before another trial starts) to elucidate if recipient mice altered their behaviour after being rewarded or not by their partners. We thus quantified the latency to go from the choice area to the reward area from the moment of the decision, as a proxy for reward anticipation (Figure 2e). Latency to access the reward area where recipients received a food pellet was significantly lower than when going to the arm where recipients did not eat (paired samples t-test: t_(11)_=-5.306, p=2.500e^-4^, BF_10_=130.755). With the same strategy as before, we calculated how much time per trial the animals spent exploring the areas close to the food receptacles (Figure 2f). We found that recipient mice spent longer time at the feeder area where they retrieve a food pellet compared to the selfish area, where they did not receive any pellet (paired samples t-test: t_(11)_=3.217, p=0.008, BF_10_=6.957). Finally, we measured the latency to return to the choice area to start a new trial from the moment the automated doors opened after reward delivery (Figure 2g). We found that recipients tended to take longer to exit from the reward area after a prosocial choice (paired samples t-test: t_(11)_=2.184, p=0.051, BF_10_=1.621).

Together, these results indicate that recipient mice displayed food-seeking behaviour by nose-poking repeatedly and almost exclusively towards the ‘prosocial’ side, and that their behaviour after a prosocial or selfish decision was very different too, being these distinct social cues that focal animals could base their decisions upon. Therefore, we next explored the behaviour of focal mice to disambiguate if they took the recipients’ actions into account for modulating their decisions.

### Focals do not modulate their behavior in prosocial and selfish trials

Beyond the average lack of preference found in the PCT, we assessed whether the behaviour of focal mice was different before and after a prosocial and a selfish choice (Figure 3a). We first measured the latency, from trial onset to choose between prosocial and selfish options (Figure 3b), where no statistically significant differences were observed (related samples Wilcoxon signed rank tests: Z=-0.471, p=0.638, BF_10_=0.334). We performed a ROI analysis to measure the time spent investigating both choice ports (Figure 3c), and focals spent similar amount of time per trial exploring the prosocial and the selfish pokes (related samples Wilcoxon signed rank tests: Z=- 0.392, p=0.695, BF_10_=0.338). We then quantified the latency to enter the reward area after performing prosocial and selfish choices, where focal animals were always rewarded but their partners were only after a prosocial choice (Figure 3d). No differences were observed (paired samples t-test: t_(11)_=-0.512, p=0.619, BF_10_=0.291). Once inside the reward areas, we measured how much time per trial the animals spent exploring the feeder area (Figure 3e), no differences were observed depending on trial type (paired samples t-test: t_(11)_=1.712, p=0.115, BF_10_=0.894). Finally, we calculated the latency to leave the reward areas after the doors opened to start a new trial (Figure 3f), where no differences were observed depending on choice type (related samples Wilcoxon signed rank tests: Z=1.255, p=0.209, BF_10_=0.604). Together these results show that decision-maker mice did not change their behaviour when deciding to provide food or not to another conspecific, prior to making the decision nor during the reward periods, despite the differences reported in the behavior of the recipients. These observations suggest that focal mice did not perceive the food-seeking cues, nor the different reward-related behaviours that recipient mice displayed, or did not make use of this information in order to guide their decision. To further investigate this, we performed an analysis on the social dynamics happening prior and after decision to determine whether decision-makers were socially attentive and interacting with recipients, and whether they could have perceived these social cues.

**Figure 3.**
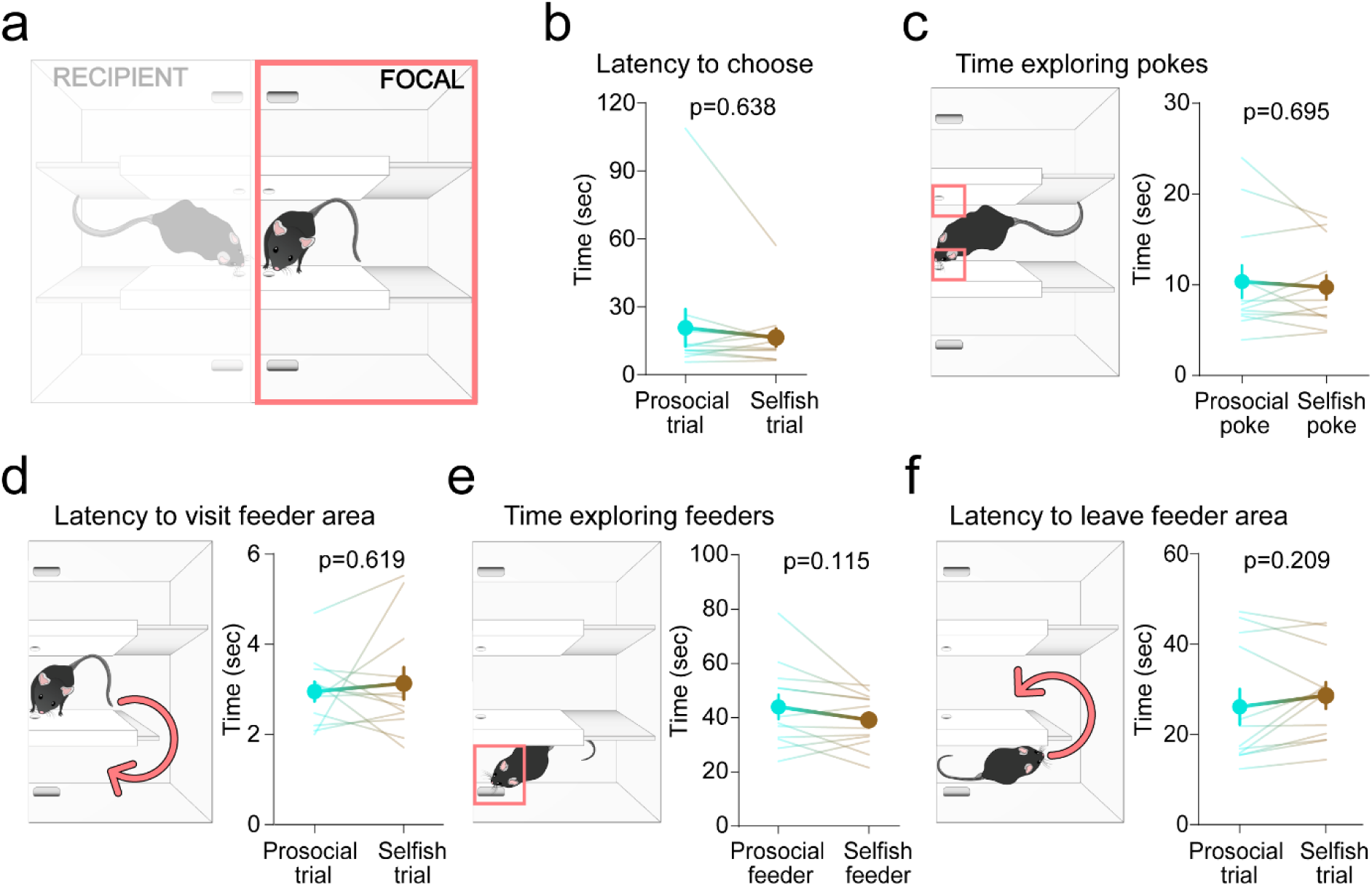
Focals do not change their behaviour according to choice type. **a**. Illustration of the arena used for the PCT. The pink rectangle indicates that the following results are focused on focals’ behavioural data. **b**. Line graph showing the latency from trial onset to choose prosocial or selfish, in seconds, where no differences were found. **c**. Time spent exploring the choice ports, measured by quantifying the frames in which the snout of the focal mice was detected in a ROI around the nose-pokes (pink squares). The time, in seconds, spent exploring both ports is similar. **d**. Latency to enter reward areas. Time in seconds from choice to detection of the focal mouse in the reward areas was not different when choosing a prosocial or a selfish choice. **e**. Time spent exploring the feeder area. Same as C for the feeder areas (pink square), in this case we also considered the detection of the head label to avoid data loss by occlusions from the walls separating the different areas. Focals do not differ on the time spent on both feeder areas. **f**. Latency to leave reward areas. The time in seconds since the automated door opens after reward, until the focal goes back into the choice area to start a new trial, is not different in prosocial or selfish trials. For all graphs: degraded thicker line shows mean±SEM, thinner lines represent data from each individual. Blue = prosocial, brown = selfish.

#### Social interactions prior to choice during the prosocial choice task

With pose-estimation data obtained from DLC we first analysed the social interactions happening from trial start to the moment of the decision to examine whether focal mice were attentive to the displays of preference performed by their recipient conspecifics (Figure 4a). We extracted the *x,y* coordinates of different selected body parts across the frames of the experimental videos and calculated different quantitative parameters that would allow the study of social dynamics of the two interacting animals. We first calculated the social (*Euclidean*) distance between the snouts of the two mice. Then, we set a threshold of 60 pixels (1,4 cm) distance to be considered a close interaction (i.e. nose-to-nose direct investigation through the diving perforated wall) and measured the time prior to choice that animals spent closely interacting in both prosocial and selfish trials (Figure 4b). No differences were observed in the duration of close social interactions prior to choice (related samples Wilcoxon signed rank tests: Z=0.863, p=0.388, BF_10_=0.511). However, relevant social interactions might have occurred at a distance. Thus, we calculated the time that both animals spent together in a defined ROI along the perforated and transparent partition that divides the arena (Figure 4c), which could provide more information about diverse social behaviours that might have happened before the decision was made. We found no significant differences in the time spent interacting near the division wall according to trial type (related samples Wilcoxon signed rank tests: Z=0.863, p=0.388, BF_10_=0.515). We then checked if they interacted in the partition at a similar distance in prosocial and in selfish trials (Figure 4d), and found that they were closer to each other prior to prosocial choices (related samples t-test: t_(11)_=-2.918, p=0.014, BF_10_=4.510). This difference could indicate higher social attention by one or the two interacting animals in prosocial trials or could just be a product of the maze configuration, where focals need to poke in the opposite side of recipients during selfish trials. Thus, we measured the distance of each animal to the wall when they were interacting inside the partition ROI (Figure 4e). We use this variable as measure to know which animal is driving a close interaction by proximity to the partition, and hence to the other animal, that is not possible to know from the Euclidean distance. We found that during the interactions near the partition, focals and recipients maintained a similar distance to the wall both in prosocial and selfish trials (RM ANOVA with trial type as within subjects factor and role as between subjects factor: F_(1,22)_=0.174, p=0.680 for trial type; F_(1,22)_=1.808e-4, p=0.989 for interaction; F_(1,22)_=0.078, p=0.783 for role. Simple main effects comparing trial type for focals: F_(1)_=0.166, p=0.692, and for recipients: F_(1)_=0.062, p=0.809. Simple main effects comparing according to the role for prosocial trials: F_(1)_=0.108, p=0.745, and for selfish trials F_(1)_=0.051, p=0.824). Yet, that both animals spent time together at a distance in the same space does not necessarily mean that they are paying attention to each other. Therefore, we measured the relative head orientation of both animals to get a proxy of visual interest during the moments prior to decision.

**Figure 4.**
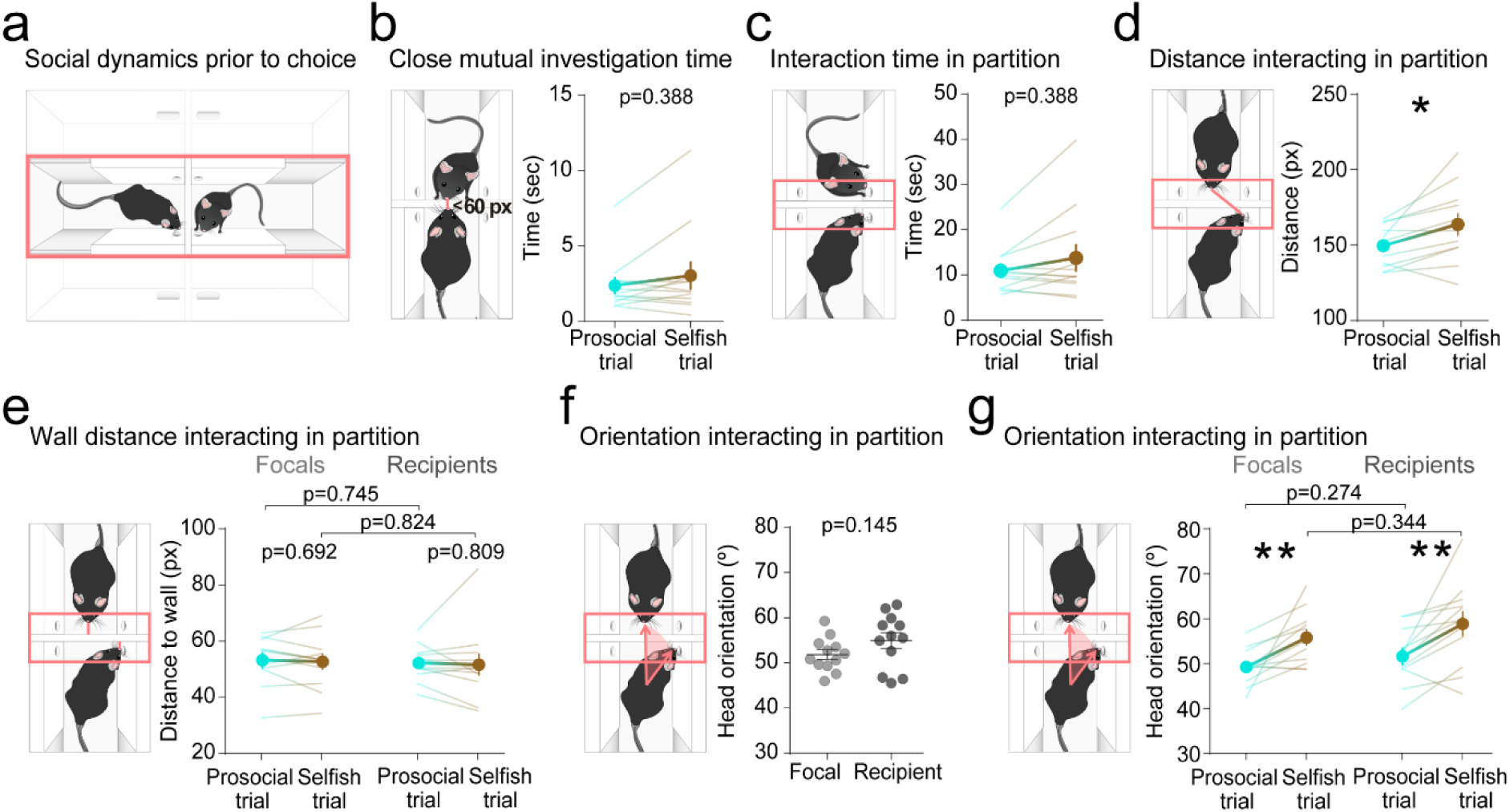
Social dynamics prior to choice. **a**. Illustration of the arena used for the PCT. The pink rectangle indicates that the following results are focused on the analysis of social behaviours in the choice area happening during the period from trial start to choice. **b**. Time in close distance. We calculated the amount of time mice interacted with a nose-to-nose distance lower than 60px (1,4 cm), which we considered to be a proximal interaction. We found that mice spend similar time interacting close to each other before a prosocial or a selfish decision. **c**. Interaction time in partition. Measurement of time spent by both mice detected together in a ROI around the divisor wall. Results show no differences on the time spent by both animals close to the partition before prosocial or selfish choices. **d**. Euclidean distance between mice during interactions in near the wall. Mice were closer during interactions prior to prosocial choices. **e.** Distance to the wall during interactions in the partition ROI. We measured the distance from each animal’s nose x coordinate to the partition. We found that both focals and recipients kept a similar distance to the wall both in prosocial and selfish trials, also when compared between them. **f**. Head-orientation during interactions in the partition for focals and recipients independent on the trial type, similar between them. **g.** Orientation in the partition according to the trial type. We found that both focals and recipients, interacted more oriented to their partner in prosocial trials, but they did not differ according to the role. For all graphs: degraded thicker line shows mean±SEM, thinner lines represent data from each individual. Blue = prosocial, brown = selfish.

This parameter represents how straight the body-head angle is with respect to the other animal’s head; values closer to 0 indicate an oriented position towards the other animal’s face, values closer to 180 indicate a head orientation opposite to the other mouse. We focused the analysis when the animals were interacting in partition ROI (Figure 4f). We found that independently of the trial type, focal and recipient mice were interacting with a similar orientation to each other (independent samples t-test, t_(22)_=- 1.512, p=0.145), that was in the range of (50-60°), enough so they could be gazing each other. We then explored whether their orientations differed according to the trial type (Figure 4g), and found that in prosocial trials both focals and recipients were more oriented to their partner (RM ANOVA with trial type as within subjects factor and role as between subjects factor: F_(1,22)_=22.004, p=1.117e-4 for trial type, F_(1,22)_=0.057, p=0.813 for interaction, and F_(1,22)_=1.446, p=0.242 for role. Simple main effects comparing trial type for focals: F_(1)_=11.587, p=0.006, and for recipients: F_(1)_=10.616, p=0.008. Simple main effects comparing according to the role for prosocial trials: F_(1)_=1.259, p=0.274, and for selfish trials: F_(1)_=0.934, p=0.344). However, differences found in this parameter according to the trial type might be explained by the task configuration. By these means, assuming that recipients are located near the prosocial port, when focals choose the selfish option by nose-poking in the opposite nose-port, the angle of both animals should increase.

Withall, we found that social interactions prior to the choice differed depending on the focals’ decisions. In prosocial trials animals interacted at a closer distance and were more oriented to each other, which should have enabled focal mice to perceive the food-seeking behaviour displayed by their partners.

#### Social interactions during reward periods in the prosocial choice task

We next examined the social dynamics that happened during the reward period (Figure 5a). To start, we measured the time that both animals spent socially interacting close to the feeder areas by calculating the number of frames that any of the head labels was detected in determined ROIs around the food-magasins (Figure 5b). Mice interacted for a longer time in the area where both the focal and the recipient receive reward compared to the selfish area (related samples Wilcoxon signed rank tests: Z=2.981, p=9.766e^-4^, BF_10_=59.45). During these interacting periods we calculated parameters such as the relative head orientation of the animals, and found that both focals and recipients were fairly oriented to each other (Figure 5c), being focals significantly more directed towards the recipients (independent samples t-test: t_(22)_=- 4.910, p=6.550e^-5^, BF_10_=290.919). We then explored how the head orientation of each mouse towards its partner was modulated by the type of trial during the interaction in the feeder areas (Figure 5d). Focals’ orientation didn’t differ according to the trial type but were more oriented than recipients both in prosocial and selfish trials. Recipients were more oriented to their focals in prosocial trials (RM ANOVA with trial type as within subjects factor and role as between subjects factor: F_(1,22)_=8.291, p=0.009 for trial type, F_(1,22)_=0.359, p=0.555 for interaction, and F_(1,22)_=22.497, p=9.830e-5 for role. Simple main effects comparing trial type for focals: F_(1)_=2.761, p=0.125, and for recipients: F_(1)_=5.717, p=0.036. Simple main effects comparing according to the role for prosocial trials: F_(1)_=10.947, p=0.003, and for selfish trials: F_(1)_=17.561, p=3.788e-4). Finally, we examined the individuals’ distance to the division wall while they were interacting in the feeder areas (Figure 5e). We found that both focals and recipients kept a similar distance towards the partition when analysed independently of the trial type (independent samples t-test: t_(22)_=0.106, p=0.917, BF_10_=0.375). This distance was not modulated by the trial type in the case of focals but recipients approached more to their focals in trials where they were not rewarded (i.e. selfish trials) (Figure 5f) (RM ANOVA with trial type as within subjects factor and role as between subjects factor: F_(1,22)_=8.774, p=0.007 for trial type, F_(1,22)_=14.080, p=0.001 for interaction, and F_(1,22)_=0.087, p=0.770 for role. Simple main effects comparing trial type for focals: F_(1)_=0.512, p=0.489, and for recipients: F_(1)_=16.213, p=0.002. Simple main effects comparing according to the role for prosocial trials: F_(1)_=2.114, p=0.160, and for selfish trials: F_(1)_=4.921, p=0.037).

**Figure 5.**
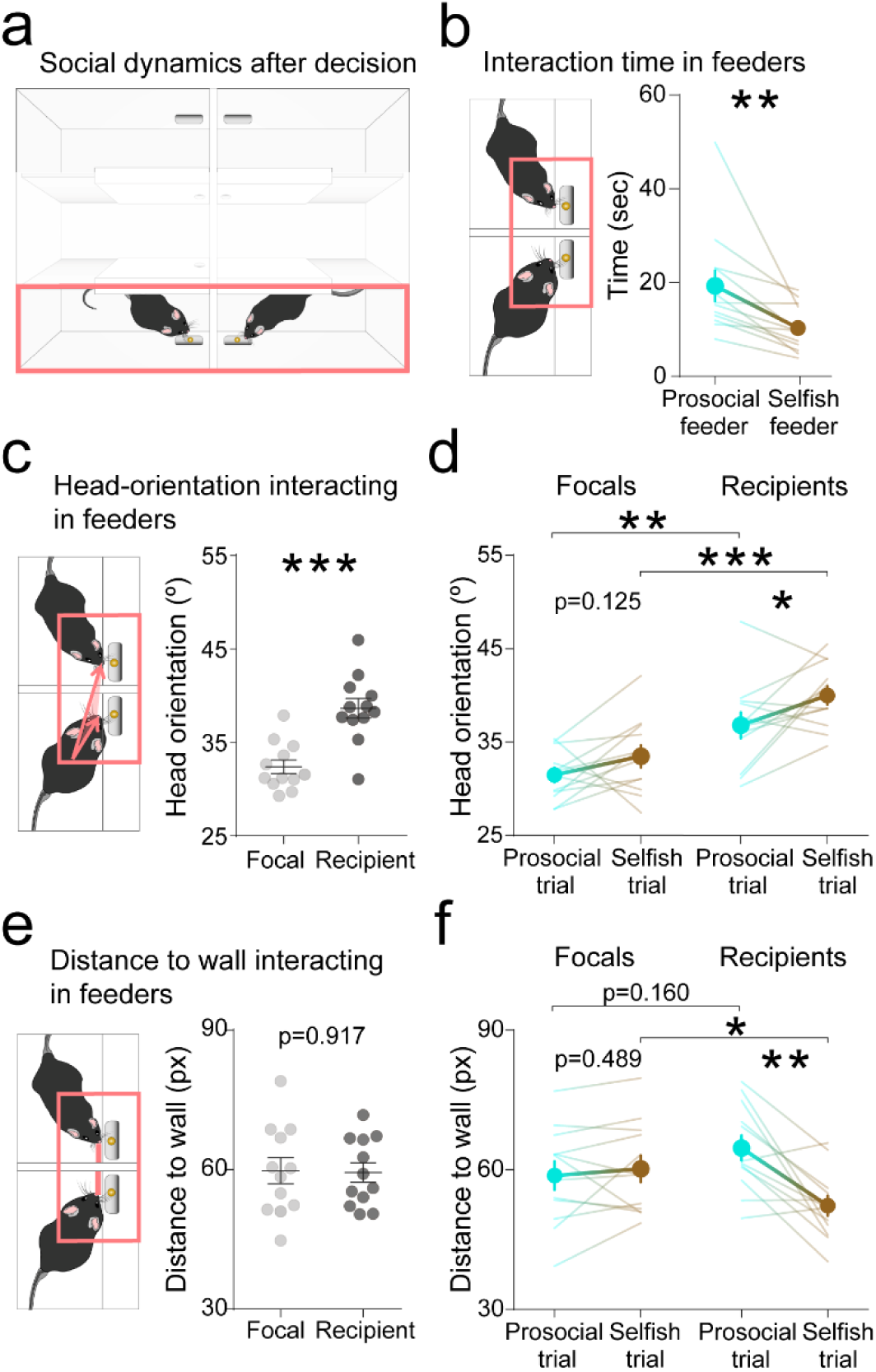
Social dynamics after decision. **a**. Illustration of the arena used for the PCT. Results are focused on the social behaviours in the reward areas during the period from choice to reward. **b**. Interaction time in prosocial and selfish reward areas (pink squares). Mice interacted significantly longer in the prosocial area. **c**. Head-orientation of focals and recipients per trial during interaction in feeder areas. Focals are more oriented towards their recipients. **d**. Head-orientation of focals and recipients during interactions in prosocial and selfish trials. Focals keep the same orientation while recipients are more oriented towards their focals in the prosocial area. **e.** Distance to partition for focals and recipients. We measured the distance to check which animal is closer to the partition, hence to the other mouse. We found no differences between focals and recipients. **f**. Distance to partition for focals and recipients in prosocial and selfish trials. Focals keep the same distance, whereas recipients are closer in selfish trials. For all graphs: degraded thicker line shows mean±SEM, thinner lines represent data from each individual. Blue = prosocial, brown = selfish.

Taken together, these results show that focal mice were oriented towards the recipient during interaction periods both prior and after the decision was made, but this social interest was not modulated depending on the choices focal animals made. Recipient animals, however, did show differences on how they interact towards their partner (i.e. orientation and distance) when focal animals decided to act prosocial or selfishly, yet none of these behaviours seemed to affect the decisions of the focals.

#### Individual differences between prosocial and selfish mice

After observing marked individual differences in focals’ prosocial biases (Figure 1e), we investigated whether their preferences would reflect differences in the behaviour of the focals, the recipients or their social interactions throughout the PCT sessions.

First, we checked that the averaged prosocial choices from the animals categorized as prosocial (n=2) or selfish (n=3) according to the permutated PCIs would actually differ, and found that, as expected, animals categorized as prosocial had a significantly higher prosocial preferences over sessions than selfish animals (Figure 6a) (Independent samples t-test: t_(3)_=11.179, p=7.672e^-4^, BF_+0_=29.270). We then found that those recipients that were paired with a prosocial focal would initially display stronger food-seeking behaviours, having a higher frequency of prosocial pokes during the first session of the PCT (Figure 6b) (Independent samples t-test: t_(3)_=3.978, p=0.014, BF_+0_=5.162), being prosocial focals more oriented to their recipients during these displays of food-seeking (Figure 6c) (Independent samples t-test: t_(3)_=8.582, p=0.003, BF_10_=9.178). Moreover, recipients paired with prosocial decision-makers tended to enter faster to the reward areas after their partners’ choices (Figure 6d) (Independent samples t-test: t_(3)_=2.923, p=0.061, BF_10_=1.831). Furthermore, when prosocial decision makers decided to behave selfishly, they displayed more proximal interactions to their unrewarded recipients (Figure 6e) (Independent samples t-test: t_(3)_=3.457, p=0.041, BF_10_=2.263). Keeping in mind the low n of animals that were not classified as unbiased, these results suggest that prosocial choices in mice are reinforced by the food-seeking behaviour of recipients and the decision-makers’ social interest to the behaviours displayed by their partners.

**Figure 6.**
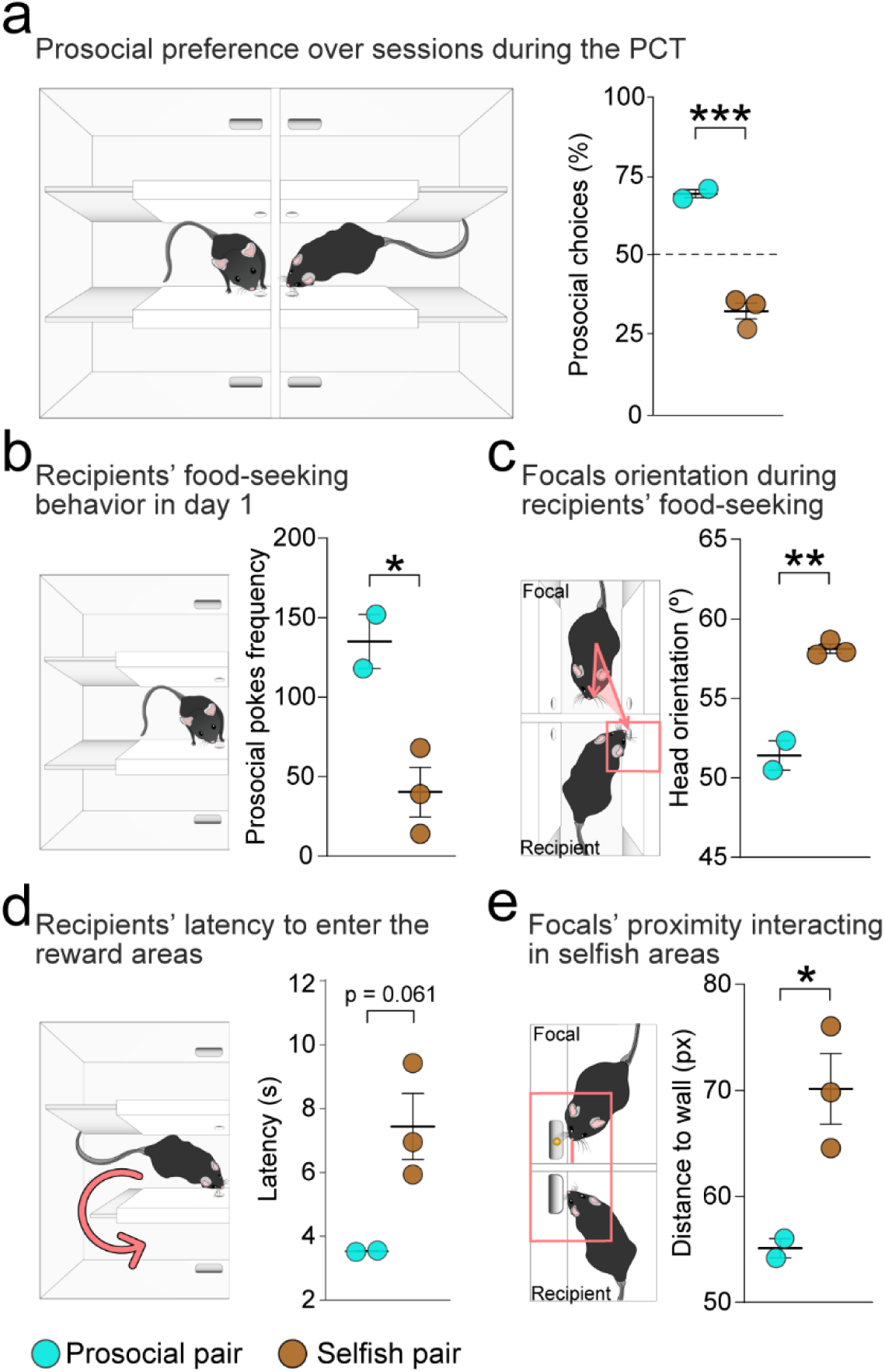
Individual differences between prosocial and selfish mice. **a**. Averaged prosocial preference over seven sessions of testing in the PCT. Prosocial focals have a higher preference of prosocial choices compared to selfish ones. **b.** Recipients’ total frequency of prosocial nose-pokes on day 1 of the PCT. We compared the total frequency of prosocial nose-pokes that recipient mice did during the first session of the PCT, according to the classification of the pairs as prosocial or selfish. We observed that those recipients from prosocial pairs do significantly more prosocial pokes compared to selfish. **c.** Focals’ head orientation during interactions when the recipient was near the prosocial nose-poke. We compared the head orientation of decision-makers when interacting with their recipients while they were displaying food-seeking behaviour prior to choice. Prosocial focals were more oriented to their partners than selfish. **d**. Recipients latency to enter reward areas after choice. Recipients from prosocial pairs entered faster to both reward areas after focals’ choices. **e.** Distance to wall for focals during selfish reward periods. We found that prosocial focals were closer to the wall during interactions in the selfish reward area.

#### Assessing prosocial choices with low-trained decision-makers

To further investigate the lack of prosociality observed in our experiments performed with mice, we considered that the individual training protocols for focal animals might have led to overtraining, making them less flexible to the new contingencies of the task in the social setting, and thus reducing the probability of adapting their behavior to the recipients’ cues. We thus evaluated whether the individual training performed in decision-makers was interfering with the emergence of prosocial tendencies in our hands. We performed an independent experiment (n=12 pairs), where decision-makers had a minimal individual training, consisting of only two sessions of fixed-ratio 1 before social testing (Supplementary Figure 2a-e), while recipients maintained their standard training schedule (Supplementary Figure 2f-h). No food-restriction protocols were used in any session of this experiment. We assessed the decision-makers’ preference and found that short training does not promote prosocial choices, as on average, focal mice had no preference for any of the options over days (repeated measures ANOVA with ‘session’ as within subjects: F_(6)_=0.909, p=0.494, BF_incl_=0.153) (Figure 7a). Interestingly, the preferences were very polarised (i.e. some focals were completely prosocial while others completely selfish) specially in the first sessions. Thus, short training seemed to promote a foraging strategy for the single choice exploitation rather than both choices exploration. Categorization in preference groups according to the Prosocial Choice Index revealed that most of the animals in this experiment (i.e. with low experience navigating in the maze) were unbiased, only one was selfish and none of them prosocial (Figure 7b, Table 1). These results indicate that, in our hands, shorter individual training did not increase the rate of prosocial choices in mice.

**Figure 7.**
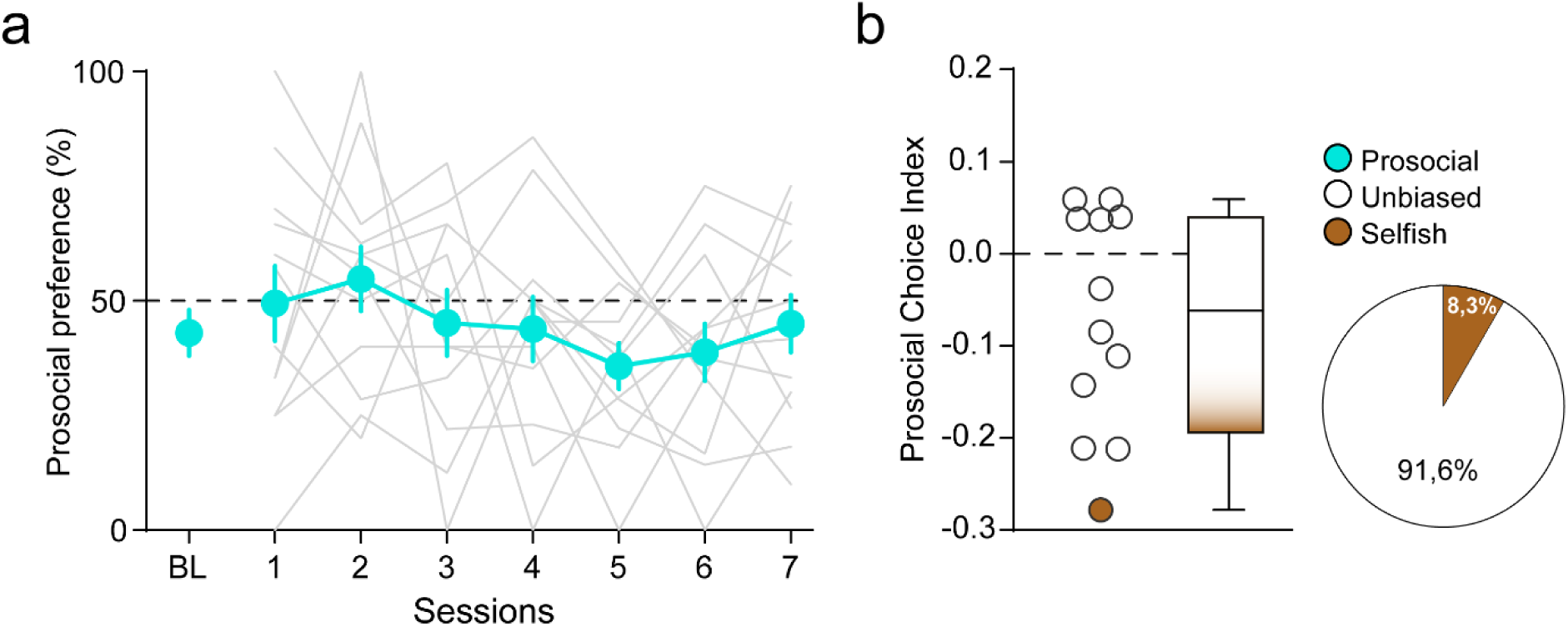
Choice preference of decision-makers with low training level. **a**. Prosocial preference of low-trained focals running the Prosocial Choice Task. Percentage of prosocial choices (Y axis) over the seven testing sessions (X axis). BL refers to baseline, used to evaluate individuals’ preference in the last two sessions of individual training. Blue thick line corresponds to mean±SEM, grey lines correspond to each individual. **b**. Distribution of Prosocial Choice Indexes. Positive values show a preference for the prosocial option, negative values indicate preference for the selfish option, and values close to 0 indicate chance preference. Blue dots correspond to prosocial mice, grey dots are unbiased and brown selfish. On the right, pie chart: distribution of mice after permutation test of Prosocial Choice Indexes.

#### Prosocial choice task with two-chamber setup configuration

So far, we evaluated prosocial tendencies in mice using a double T-maze configuration, where choice and reward delivery were separated spatially and temporally in different compartments. Next, we developed a simpler two-chamber paradigm and assessed if task configuration had an impact in laboratory mice prosociality. The design of this new setup consisted in an acrylic box, with a transparent and perforated partition in the middle which allowed mice to see, hear, smell, and partially touch each other. There were two contiguous areas, one for the decision-maker (focal mouse) and another for the recipient mouse (where delivery of reward depends on the focal’s choices) (Figure 8a). The choice ports (i.e. prosocial and selfish) were located equidistant from the recipient (to avoid possible baseline preferences for the one closer to the partner or local enhancement effects in that poke) and the recipients had one nose-port to display food-seeking behaviour. More

**Figure 8.**
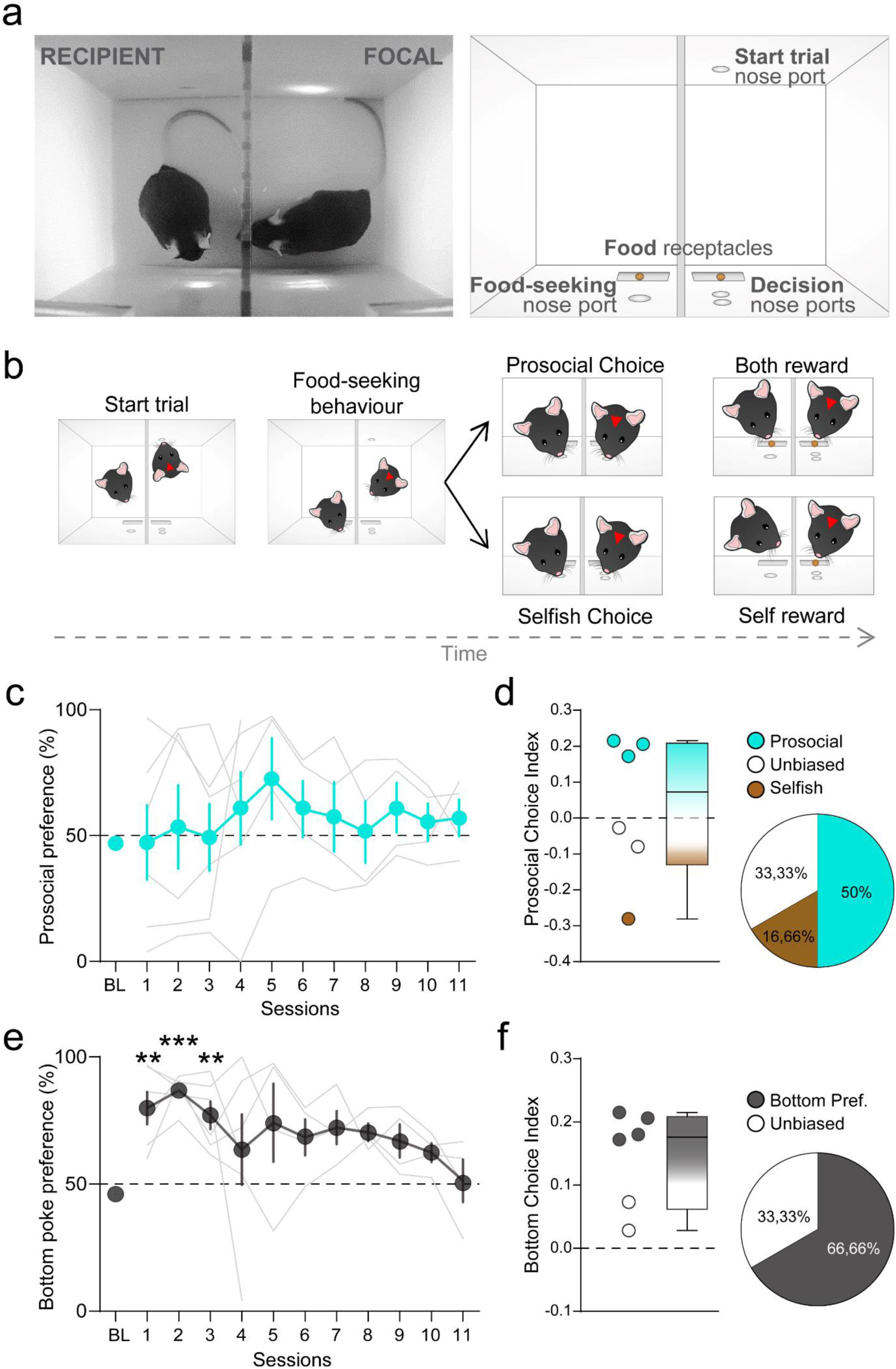
Mice prosocial choices in the two-chamber PCT. **a**. Left, real top-view image of the two-chamber setup used for evaluating prosocial decision-making with mice. Focal (on the right) is checking the recipient mouse (on the left) through the perforated and transparent partition that divides the two chambers. Right, schematic of the peripherals used inside the setup. In the focals’ side there is a nose-port to start the trials, on the opposite wall there are two other nose-ports placed vertically used for decision and below these there is a food magazine where food pellets are delivered. In the recipients’ side there is a single nose-port to display food-seeking behaviour and below there is the food magazine for rewards. **b**. Timeline structure of the prosocial choice task. Trials begin when the decision-maker pokes into the nose-port to start, then the recipient mouse needs to poke into the nose-port to display food-seeking behaviour which activates the pokes for decision of the focal mouse. Poking into the prosocial port will deliver a food pellet to both animals however, choosing the selfish port will reward only the focal mouse. **c**. Prosocial preference of mice in two-chamber arena. Percentage of prosocial choices made by all focal mice (Y axis) during 11 experimental sessions on consecutive days (X axis). BL referrers to baseline, used to assess the preference of the focals on the last two days of individual training. Blue line corresponds to mean±SEM, grey lines correspond to data of each individual. At population level, prosocial preference was not different from chance (50%) in any experimental day. **d**. Distribution of Prosocial Choice Indexes. Positive values show a preference for the prosocial option, negative values indicate preference for the selfish option, and values close to 0 indicate chance preference. Blue dots correspond to prosocial mice, white dots are unbiased and brown selfish. On the right, pie chart: distribution of mice after permutation test of the Prosocial Choice Indexes. **e**. Preference for bottom choice. Percentage of choices towards the bottom option (Y axis) over the testing sessions (X axis). Black line corresponds to mean data from all individuals, grey lines correspond to each individual data. There is a significant preference for the bottom option already in the first 3 days, which slowly decays over sessions. **f**. Distribution of bottom choices. Positive values show a preference for the bottom option, negative values indicate preference for the upper option, and values close to 0 indicate chance preference. Grey dots represent mice with preference for the bottom option, and white represent unbiased. On the right, pie chart: distribution of mice after permutation test of the bottom choice index in percentages.

During individual training of the decision makers, focal mice were trained to poke into both options, so by the end of the individual training they showed no strong preference for any of the options (Supplementary Figure 3b) (one sample t-test against chance (50), t_(5)_=-1.324, p=0.243, BF_10_=0.698). Once the individual training was fulfilled, mice underwent the prosocial choice task (PCT) (Figure 8b, Supplementary Video 2). We tested six pairs of cage-mate male mice (C57BL/6) during 11 sessions of 40 minutes. As food deprivation has been previously reported to increase prosocial tendencies in a similar two-chamber paradigm (Scheggia et al., 2022), for this set of experiments, we mildly food restricted the animals before undergoing the PCT, to increase their motivation for obtaining the food pellets. Under this schedule, we evaluated if mice develop a preference towards the prosocial option and found that mice did not have any consistent preference over sessions in this double-chamber paradigm (repeated measures ANOVA with ‘session’ as within subjects factor: F_(10)_=1.105, p=0.39, BF_incl_=0.335) (Figure 8c). Yet, we observed marked individual differences over sessions, already present on the first session of the PCT (Figure 8c, grey lines).

Indeed, with this double chamber paradigm, the categorization of focal animals according to their PCI scores after the permutation test revealed that 50% of the animals were prosocial, 2 focals were unbiased and only one was selfish (Figure 8d, Table 1), which represents a much higher proportion of animals compared to our double maze configuration paradigm. However, these preferences for the prosocial or selfish option were explained by a general preference for the poke that was placed closer to the food magasin, which did not required the animals to rear, and thus was less costly (repeated measures ANOVA, with ‘session’ as within subjects factor: F_(10)_=2.42, p=0.03, BF_incl_=2.6) (Figure 8e), especially during the first three sessions of the PCT, when they are learning the new contingencies of the task: (one-sample t test against chance (50%): t_(5)_ = 4.697, p=0.005, BF_10_=11.069 for session 1; t_(5)_=14.185, p=3.136e-5, BF_10_=600 for session 2; t_(5)_ =4.921, p=0.004, BF_10_=12.86 for session 3). We also calculated the PCI for the bottom preference to assess individuals’ differences in the preference, and found a significant increased preference from chance (one sample t-test against 0: t_(5)_=4.65, p=0.003, BF_+0_= 21.29) (Figure 8f, Table 1). These results suggest that the marked individual differences we observed in the prosocial choice task were explained by a bias towards the option that would require less effort and was not related to prosocial biases.

## Discussion

In this work, we introduced a new custom-made behavioural paradigm for the study of prosocial choices in mice. The design and development of this setup was based on the one previously described in (Márquez et al., 2015) to assess prosociality by reward provision in rats. Three main reasons motivated the adaptation of this paradigm to mice: **(1)** the scarcity of studies for assessing reward-based prosociality in mice **(2)** the use of the larger genetic toolbox available nowadays for the manipulation and monitoring of neural circuits for mice in comparison to rats, **(3)** the possibility for direct interspecies comparison using the same paradigm. To date, different labs have developed and used their own paradigm to give answers to similar questions about prosociality. On the one hand, being able to develop a specific paradigm to flexibly address a question of interest is very relevant and beneficial to further expand our knowledge. However, on the other hand, this might be problematic when obtaining conflicting results between studies that use different training or testing procedures, which makes it difficult to interpret the results due to the variability in the different factors included among them. Thus, replication of results using the same paradigms and protocols would be useful and would allow for direct interspecies comparison.

Withall, there are diverse PCT paradigms used to assess reward provision in rats that, in general, find these to be prosocial by preferring the choices that reward a conspecific (Gachomba, Esteve-Agraz et al., 2024). In contrast, the results found in the few mice studies give contradictory conclusions, which might arise from the clear differences in the paradigms and testing protocols used. To our knowledge, only two studies have tested mice prosociality using reward-provision paradigms (Misiołek et al., 2023; Scheggia et al., 2022). Both studies used double-chamber arenas which separated the animals through a transparent and perforated partition, however in Misiolek’s work, focal animals had to choose alone in a compartment and only had ‘access’ to recipients’ cues during the reward, once the decision was made. In Scheggia’s study, choice and reward consumption happened in the same compartment, and mice were always accessible to each other. They further tested how different partitions affected prosocial choices finding that a metal mesh (i.e. the only allowing for social contacts) promoted prosocial choices. Another main difference between these paradigms is regarding the trial structure during testing. In Misiolek’s study, mice performed 4 testing sessions with 6 forced trials and 16 free choice trials each, with limited time per trial. On the other hand, Scheggia tested mice for 5 sessions in which they could perform *ad libitum* choices for 40 minutes. In our experiments, animals always had access for social interactions prior and after choice and could perform *ad libitum* choices for 30-40 minutes during 7-11 sessions (i.e. in the double-T and double-chamber paradigms respectively).

Despite the clear differences in the paradigms and testing protocols used, a main difference found between the studies is concerning prosocial reward provision in mice to be sex-dependent in two opposing directions. While (Scheggia et al., 2022) found that most of tested male mice (75%) were prosocial, in comparison to only 47% of tested female mice. (Misiołek et al., 2023) found that female mice are more willing to choose the prosocial choice more often than males (75% prosocial female to 30% prosocial male mice). Our results using our double T-maze seem to favour this second study, as only 17% of the tested males were prosocial (in the standard training protocol). However, we limited our experiments to test only male mice, as we first aimed to validate the task and paradigm, which for rats provides evidence for no sex differences (Gachomba et al., 2022).

In an attempt to explain the generalized lack of prosociality found in our double-T maze experiments with mice, we performed a detailed quantification and analysis of social and individual behaviours occurring during this task. We first focused on the recipients’ actions and found that they displayed robust food-seeking behaviour that was almost exclusive towards the prosocial nose-port. In previous work, we found these actions to be important and necessary for the emergence of prosocial preferences in rats, as when prevented, decision-maker rats did not become prosocial (Márquez et al., 2015). Additionally, we explored several behavioural measures from the recipient animals prior and after the decision-makers’ choices. We found that recipients displayed very distinct behavioural patterns in prosocial and selfish trials, reflecting that they were aware of when they were going to be rewarded and showed clear behaviours that decision-makers could have used to guide their decisions. However, despite the clear behavioural cues from recipients, we found that focal animals behaved similarly whether they decided the prosocial or the selfish option. Because decision-makers did not seem to be affected by the different behaviors displayed by recipients depending on the trial type, we asked whether animals were actually socially interacting and attentive to the behavior of their partners. In broad terms, we did not find any predictive variable in their social interactions prior to choice, although the social distance and orientation they displayed should have enabled focals to perceive the food-seeking and reward-related behaviours their partners displayed. After choice, those differences found in their social dynamics arose from the behaviour of recipients according to the trial type, whereas focals remained invariant.

When exploring the individual differences observed in our mice, and considering the low sample of the two groups identified according to their choice preference (prosocial, n=2; selfish, n=3), results show that prosociality in mice and rats might be regulated by similar proximate mechanisms. Indeed, we found that recipients of those few prosocial decision-makers were better a displaying food-seeking on the first session, which as mentioned previously, is a necessary action for focal rats to develop a prosocial preference (Gachomba et al., 2022; Márquez et al., 2015), and they entered faster to the reward areas after all focals’ choices. We also noted that prosocial decision-maker mice oriented to their recipients during their displays of food-seeking behaviour more than selfish ones, and were closer to the partition during social interactions in the selfish reward area, suggesting social interest to the lack of reward to their cage-mates. Altogether, these insights suggest that behaviours and social interactions in specific moments (food-seeking during the first session and interactions in reward areas) were especially relevant for prosocial focal mice to pay increased attention to their partners’ behaviour, which in turn probably eased the understanding of the reward contingencies of their choices.

To keep further investigating the general lack of prosociality observed in our experiments performed with mice, we considered the individual training protocols used as a possible factor interfering with the decision-making process during the social task. It has been suggested that prolonged training of an instrumental action like nose-poking, can make such behaviour become habitual and thus, less flexible and goal directed (Thrailkill & Daniels, 2024). Furthermore, prior individual training has been demonstrated to have an effect in prosocial actions to avoid harm to others in rats (Hernandez-Lallement et al., 2020). Thus, the invariant and comparable individual behaviours displayed by focal mice in prosocial and selfish trials together with their average lack of preference found in these experiments could reflect choosing both options similarly as a habitual action, thus becoming less dependent on the actions and behaviours displayed by their partners. In the work of Scheggia and colleagues, decision-maker mice learnt the task contingencies during social testing and most male mice ended up being prosocial. Additionally, they assessed whether sharing food with a recipient could motivate a change in choice preference when having already a prior stable preference alone. They found in this case that most mice also switch their preferred option to favour the partner, although with a weaker effect (Scheggia et al., 2022). This idea suggests that mice tested in our paradigm could show prosocial tendencies with a different (lower experience) individual training schedule for the decision-makers. Short training of instrumental actions prior to the PCT did not favour prosocial choices, and in any case, diluted the individuals’ preference by promoting a foraging strategy for the single choice exploitation rather than promoting their natural tendency to alternate between choices (Lalonde, 2002). Overall, mice showed very polarized choices from one session to another, revealing that no decision-maker was prosocial over sessions and most remain unbiased (except for a selfish one).

Although in our hands, mice prosocial choices for reward provision were not very widespread, these experiments served to compare those results with rats, who naturally display a prosocial preference in different settings. We considered that our double-T paradigm could either be too cognitively demanding for mice or did not fully capture the ability of these animals to display prosocial choices in this context. Thus, in a final attempt to probe mice prosociality in a reward-based context, we developed a similar task with a simpler layout (i.e. two-chamber PCT paradigm), more similar to the one from Scheggia et al., 2022, but with main differences. **(1)** the recipient mouse had an active role in the task, by displaying food-seeking behaviour in a single nose-poke to incentivize focals’ decisions. **(2)** the choice nose-pokes of the focal mouse were placed vertically and equidistant to the recipient, instead of horizontally, in order to avoid possible local enhancement effects towards the option being closer to the recipient. **(3)** all animals underwent an individual training prior to social testing. The results from this experiment reflect that this setup configuration was not optimal for assessing prosocial choices in mice. By the end of the individual training, focal mice displayed no bias for any of the two options, however during social testing (and already in the first session) most of them had an overall preference for the nose-port placed at the bottom, requiring a lower effort. In this way, we argue that when the reward contingencies of the task changed during social testing, decision-makers could possibly not disentangle the different social cues that, in this case, happen in the same area and close in time, and which result important for the emergence of prosocial behaviours in rats. Thus, in our hands, a simpler layout of the PCT did not better capture the ability of mice to display prosocial choices and actually hindered the interpretation of their choices.

All things considered, in our experiments, prosocial tendencies in reward provision in mice are not as widespread as for rats and come with marked individual differences. In those few animals we found to be prosocial in the double-T paradigm, the behavioural mechanisms promoting prosociality are similar to the ones in rats. These results provide a framework to perform direct interspecies comparisons, thus once we understand the neural circuits underlying these intricate behaviours in rats, performing gain of function experiments with mice could bring about very promising results.

### Mice and rats are different in the wild and in the lab

Rats and mice are widely used models for studying mechanisms of mammalian social behaviour and cognition, and while some might think that a rat is just a “bigger” mouse, substantial differences in social cognition and behaviour have been described between these two species, likely linked to their distinct natural social structures (Ellenbroek & Youn, 2016). It has been described that both rats and mice live in hierarchical groups in the wild, yet rats are much less territorial and the hierarchy between males is not as absolute as for mice (Schweinfurth, 2020). Rats are commonly found to live in mixed-sex colonies, where all males mate with all females, and they have further been described to forage for food together (Weiss et al., 2017). In contrast, mice are commonly found to live in more territorial structures, which are founded by a single male that mates with all the females (Lipp & Wolfer, 2013). As a result, interactions between males are much rarer, and when they occur, are more aggressive and territorial in nature. These differences in the natural social dynamics in the wild also translate when studying them in laboratory settings. For instance, different studies have examined the motivation for seeking social contacts in both rats and mice. Using a socially-conditioned place preference test, (Kummer et al., 2014) described that most of the rats found the socially-paired compartment rewarding, spending more time, compared to half of the tested mice, even describing that a small proportion could find it aversive. Moreover, rats found social interaction as rewarding as 15mg/kg body weight cocaine, whereas mice preferred this dose much more than social interaction. In this line, others have described similar effects comparing the motivation for seeking food or social contacts between rats and mice. (Reppucci & Veenema, 2020) developed a social vs. food preference test, and found that in sated conditions, rats were generally more social-preferring whilst mice were unbiased (Reppucci et al., 2020). In food deprivation conditions, their preferences switched; rats had no bias whereas mice preferred the food stimulus. (Netser et al., 2020) replicated such observations and further examined the dynamics of their social investigation. The authors found distinct exploration strategies employed by rats and mice when choosing to interact with familiar or unfamiliar conspecifics. Their results demonstrate a stronger immediate motivation for interactions with same-sex social stimuli in male rats compared to mice. Moreover, species-specific characteristics also bring about other differences in addictive and impulsive behaviours and cognition (*for review see* (Ellenbroek & Youn, 2016)), but also in emotional processing and transfer. Both rodent species can be affected by and respond to negative affective states of conspecifics at comparable levels (Hernandez-Lallement et al., 2022). Yet, diverse factors affect them differently: familiarity (longer time living together promotes stronger emotional contagion in mice, but lower in rats), strain (modulates emotional contagion in mice but not in rats) or prior experience with emotional stimuli (pre-exposure in rats can double the emotional contagion response while no effect is found in mice) (for review see (Hernandez-Lallement et al., 2022; Keysers et al., 2022)).

Thus, one could argue that the conflicting results found in the two aforementioned studies assessing prosocial choices in mice could arise from any of these components that differ in the natural repertoires displayed by rats and mice. From an ecological standpoint, these two rodent species are considered prey animals, but mice more so than rats, as the latter can often be regarded as predators of mice (Campos, et al., 2013; Liu, et al., 2016). Furthermore, sharing food with other individuals always reduces the amount of potential food available to oneself (Chang et al., 2011). Considering the differences found in the natural social dynamics of mice and rats, it would not be surprising that prosociality measured in reward provision paradigms differed substantially between these rodent species, favouring the appearance of prosocial choices in rats, in contrast to the more solitary and competitive nature of male mice. Yet, mice social behaviours are present, and even prosocial choices can be found, but might not be prioritized as a reflection of their ecological dynamics. Furthermore, different levels of sensitivity to the affective states of others could represent a causal factor for the differences observed in reward-provision. Both Misiolek and Scheggia studies tested for affective state discrimination, in negative or relieved conditions, the former finding similar levels between male and female mice, and the latter finding a link between emotional contagion and increased prosocial choices. Their results contribute to the growing body of work finding that emotional contagion and detection of negative affective states of others in mice are robust responses. However, the generalized lack of prosociality found in our work and the contradictory evidence of these two other studies could reflect that mice are not so consistent in paying attention to the reward of others. Finding further evidence about a sensitivity to positive affective states of others in mice is fundamental to fully understand their range of emotional processing abilities.

Finally, mice often need substantially longer training and habituation sessions to perform certain tasks that rats do (Jaramillo & Zador, 2014), and additionally experience more stress and anxiety (Ellenbroek & Youn, 2016), probably reflecting their prey nature. Moreover, the literature about prosocial behaviour in rodents shows an inequal distribution of mice and rats used in prosocial studies (Gachomba, Esteve-Agraz et al., 2024), which might be related to the cognitive demands of the tasks. Thus, those tasks based on instrumental actions to measure prosocial behaviours use mostly rats (namely *rescue, reward provision and harm aversion* paradigms). In contrast, mice and rats are comparably used in consolation paradigms, which measure immediate natural responses that require no learning to be displayed. All of these species-specific variations we have been discussing emphasise the importance of choosing an appropriate animal model for studies in behavioural neuroscience, especially for those with a social scope.

## Conclusions

In this work, we aimed to study prosocial choices in reward provision paradigms in mice that, as reviewed in the literature (Gachomba, Esteve-Agraz et al., 2024), is one of the least investigated. Providing extra evidence in this direction is essential to unravel the conflicting results obtained by the few studies on reward provision in mice. Overall, the results of the current work have revealed interesting insights into behavioural substrates of prosocial choices in mice. In our hands, C57BL/6J male mice do not show prosocial tendencies at the population level, or at least not as widespread as for rats. Interestingly, we find that there are marked individual differences in prosocial choices, and those mice who develop a prosocial preference are those paying more attention to the recipients’ behaviour during reward delivery, or those whose recipients more clearly display its food-seeking behaviours. We highlight here that these are factors that we previously demonstrated to promote prosociality in rats, suggesting that prosocial tendencies are not overly prevalent in laboratory mice, but when they emerge seem to depend on behavioural elements similar as for rats.

## Materials and Methods

### 1.1. Animal subjects

**Mice**: 60 adult male C57BL6/J mice (632C57BL/6J, Charles River, France) were used, aged between 49-55 days at arrival to our facilities, with a body weight of 25±2 gr. Upon arrival from the commercial vendor, mice were group-housed (4 mice per cage) and maintained with *ad libitum* access to food and water in a reversed light cycle (12 hours dark/light; lights off 8 AM), in controlled temperature conditions. Paperboard and transparent acrylic cylinders were used as environmental enrichment in the home-cage. Mice were left undisturbed in their home-cages for the first two weeks at our Animal Facility to allow them to reverse their circadian rhythm and acclimate to the new environment and routines. Body weight was controlled weekly. All procedures were carried out following the European Commission guidelines (2010/63/EC).

### 1.2. Experimental procedures to investigate prosocial choices in mice

We developed two different setups to evaluate prosocial tendencies in food foraging contexts in mice, one with a T-maze configuration (Akam et al., 2022), and a second one with a double chamber configuration. Both apparatuses were fully automated, in order to minimise interference by the experimenter while at the same time allowing for a precise control and detailed monitoring of the behaviour of the interacting individuals. In both tasks, the choices of an animal (*focal*, the decision-maker in the task) determined reward delivery for the *recipient* partner, allowing preference for ‘prosocial’ vs ‘selfish’ choices to be examined over sessions. Focal animals reported their choices by nose-poking between two available nose-pokes: one that provided food for itself and the recipient animal (prosocial choice) and another one that only rewarded itself (selfish choice). Recipient animals displayed attempts to obtain the reward by nose-poking repeatedly into a single nose-port. Mice worked for palatable pellets (20mg Dustless Precision Pellets, Bioserv #F0071) that were automatically delivered by a custom-made pellet dispenser into a food-receptacle. Before running any procedure with the animals, mice were habituated during a week to the experimenter by handling them for 5-10 minutes per day and to the food pellets used in the experiments to avoid stress related to neophobia.

The setups were derived and adapted from the Prosocial Choice Task (PCT) developed for rats by (Márquez et al., 2015), where two different processes were identified as crucial for the emergence of prosocial decision-making: (1) the food seeking behaviour displayed by the recipients of help while trying to obtain the food prior to the focal’s choice, and (2) different reward contingencies in the reward areas of each choice and putative social information exchanged during these moments. The tasks developed and presented in this thesis were designed to include these two processes for the study of prosocial choices in mice. The differences in the structure and configuration of the two setups are explained below.

#### 1.2.1. Maze-based configuration for prosocial decision-making task

The behavioural setup consisted of a fully automated double T-maze (Figure 1a-b). Each T-maze consisted of a central corridor (choice area) with nose-poke ports on each side and two side arms (reward areas) each with a food receptacle connected to a pellet dispenser at the end. Access from the central choice area to the side arms was controlled by custom-made automated pneumatic sliding doors. Each individual maze (15 x 22 x 15 cm) was built with laser cut white acrylic (3 mm) and was connected to the other by a transparent and perforated acrylic (3 mm). This transparent partition in the middle of the apparatus divided the maze in two, one for the decision maker and another for the recipient of help. For each individual maze, the central choice area was separated from the lateral reward areas with transparent acrylic walls, which allowed visibility of the animal in the side arms of the maze enabling tracking in the entire maze with one camera placed above the setup. These walls (12 mm) contained the mechanisms for the sliding doors (made from 3 mm transparent acrylic), animal’s position IR detectors and 3D printed nose-pokes with sensors. All inner walls from the maze were gently scuffed with a fine sandpaper to avoid reflections of the mice in the walls that could interfere with automated pose estimation of the animals.

The task comprised two separate stages: (1) Individual training; in which animals learnt to navigate in the maze individually, opened doors by poking the ports in the central arms and retrieved pellets in the side arms. (2) Social task; where the decisions of the focal animal controlled the doors in both mazes, and determined rewards for both itself and the recipient animal.

##### 1.2.1.1. Individual training protocols

During individual training, all animals were first habituated to the individual T-mazes in two sessions of 15 minutes, in which free exploration of the arena was allowed (i.e., all doors were open and nose-pokes and infrared beam detectors were inactive). Several food pellets were available for consumption in the food-magazines and floor of the maze in order to habituate animals to them. During the second session, mice were habituated to the gating of the automated doors, by opening and closing them non contingently of the animals’ behaviour.

Then, mice were trained for three days to poke in the nose ports under a fixed ratio 1 (FR1) schedule (i.e. one nose-poke into the cued port required for obtaining a reward) in order to open the door that gave access to the food magazine where a food pellet was delivered for consumption. Both pokes were active during this stage and both sides were rewarded. Mild food restriction was performed during this early training, by removing the home-cage food 2 h prior behavioural testing. Once training sessions finished, mice were allowed to eat *ad libitum* for the rest of the day.

Mice were randomly assigned to be the decision-maker (focal) or recipient of the help and tested in the social task with one of its cage-mates. From this moment onwards, focal and recipient mice were trained differently, and their roles were fixed throughout the entire experiment.

###### 1.2.1.1.1. Focals’ standard individual training

Focal animals continued individual training under a FR1 schedule during ten days for 20-30 minutes, where side biases were evaluated (Supplementary Figure 1a). Briefly, a single poke in either of the two available LED-cued ports in the choice area triggered the opening of that same side door, allowing access to the lateral arm from where the animal could retrieve a pellet in the food magazine. An IR detector allowed to identify the moment when animals had reached the feeder area at the end of the lateral arm, moment in which the door safely closed in their back. Animals were allowed to retrieve and consume the pellets for a period of 10 seconds. Then, the door of the reward area opened again, allowing mice to go back to the choice area to start a new trial. Focal mice were allowed to freely choose any of the two pokes and hence get rewarded in the corresponding reward area.

###### 1.2.1.1.2. Focals’ reduced individual training

A shorter individual training was designed to assess whether the standard one was making animals to be less goal directed and inflexible in their choices once tested in the social task (*see results section* **Error! Reference source not found.**, Figure 7 and Supplementary Figure 2). In brief, focal animals in this experiment went through the two initial sessions of maze habituation. Then, they only performed a single session under fix ratio 1 schedule at the beginning of the training protocol, and a short training session just prior to social testing. This last session was limited to 20 mins or 6 trials, whatever was reached first. These two sessions were considered as the baseline for the side preference used to compare the prosocial preference during the PCT.

###### 1.2.1.1.3. Recipients’ individual training

During individual training for the recipient animals (Supplementary Figure 1d), only one of the nose-poke ports in the central area was active (randomly assigned to be the right or left poke) cued with a LED, and the number of pokes required to access the reward arm increased over the training sessions. The rationale of this protocol was for recipients (1) to show a clear preference for only the rewarded side of the maze in the social task and (2) to actively display food-seeking behaviour (nose-poking repeatedly). The first training sessions started with a FR1 schedule (i.e., only one nose-poke was necessary to open the door giving access to the reward). The quantity of pokes necessary to access the reward increased up to FR6 according to individual performance of the animals, thus ratio increased automatically when a given animal did five successful trials in a given FR schedule within a session (i.e., poking the required number of times, with a delay of less than 2 secs between pokes). Then, recipients were further trained under a variable ratio five schedule (VR5: pseudorandom list of pokes frequency needed to open the door, with an average of 5 pokes). In the last two sessions, recipient mice performing under this VR5 were forced to visit the unrewarded arm in 10% and 20% of the trials, in order to habituate them to enter and exit from the unrewarded area to initiate another trial. Finally, from the second day of PCT, a brief individual training was performed to the recipients before social testing to avoid extinction of food-seeking behaviour, as during the PCT they were not in control of their own reward delivery anymore.

##### 1.2.1.2. Prosocial Choice Task protocol

During social testing, a pair of animals (focal and recipient) from the same home-cage were placed in the double T-maze, one in each side of the maze, separated by a transparent perforated partition. In the social task, although both mice had access to the nose ports of their corresponding mazes, only those of the focal were active, and these controlled the automated doors of both mazes (i.e., a single poke to either port made by the focal animal opened the corresponding side doors in both mazes). The trial started when both mice were in the choice area. Recipient animals displayed food-seeking behaviour (poking into the port on the side where it would receive reward: prosocial side) while the focal animal controlled the recipient’s access to the food-baited arms. Importantly, the focal animal was rewarded for accessing either side, while the recipient animal was rewarded only on one side. The choice made by the focal animal therefore, determined whether the recipient animal received reward or not. Prosocial choices referred to choosing the side of the maze that provided access to food for both animals, whereas selfish choices referred to choosing the side of the maze that only provided food to the focal and not to the recipient. In this way, both choices provided the same amount of reward to focal mice. The reward of the focal animal was available immediately in the food receptacle after the choice was made, however, the recipient mouse only received its pellet once both animals were in the reward area, ensuring that information about the recipient receiving or not the food was available for the focal animal in each trial. Ten seconds after both animals entered the reward area, the doors of the recipient animal opened allowing it to return to the choice area and, once detected there by the corresponding infrared beam input, the focal’s door opened allowing it to go back to the choice area to initiate a new trial. Seven sessions of 30 minutes were performed for each pair of animals. Animals were left to feed *ad libitum* during the entire period of social testing.

#### 1.2.2. Two-chamber configuration for prosocial choice task

This behavioural setup consisted of a custom-made white acrylic arena (16 cm long x 10 cm wide x 15cm high) which was divided in two individual chambers, one for each animal of a pair (8 cm long x 10 cm wide x 15 cm high). The two chambers were separated by a perforated and transparent partition that allowed the exchange of multimodal sensory information (Figure 7a-b). In each chamber there was a food-magasin connected by a tube to a custom-made pellet dispenser. In the decision-maker chamber there was a vertical double nose-port that animals used to display their choices, placed above the food receptacle. A start trial port was placed on the opposite wall. In the chamber of the recipient of help animal, there was a single nose-port above the food-magasin.

##### 1.2.2.1. Individual training protocols in the two-chamber prosocial task

Before undergoing social testing, mice went through individual training according to their future role in the PCT. To this purpose we used a single chamber arena (8x10x15cm white acrylic box), which mimicked one chamber of the social setup, with a food receptacle and a dismountable poke-wall that allowed different training for focals and recipients. During the first two sessions, all mice were placed alone in the chamber to allow free exploration and habituation to the arena for 10 minutes. Habituation to food-pellets was obtained by allowing animals to freely consume the available pellets placed on the floor of the chamber and in the food-receptacle. Then, all mice followed an individual training session with FR1 schedule for obtaining rewards, during three sessions, after which we randomly assigned the roles for the future social task (2 focals – 2 recipients per homecage). Individual training from this point diverged for focals and recipients. Short food restrictions (2h before the behavioural testing) were performed in all stages of the individual training to increase motivation for food-seeking behaviours.

###### 1.2.2.1.1. Focals’ individual training

Decision-maker mice were trained to obtain rewards under a FR1 throughout all the individual training (Supplementary Figure 3a). Because of the configuration of this setup, focal mice reported their willingness to start a trial in a self-paced manner, by performing a poke in the start trial poke, after the 10th individual training session. During the early sessions of individual training, some of the focal mice showed an increased preference for the bottom poke (the one closer to the food magasin and that did not require to rear). When strong biases were observed, the preferred poke was blocked for some trials (i.e. pokes were not followed by reward delivery) forcing animals to explore the non-preferred option. Baseline sessions before the social task did not contain forced trials and thus reflected the individual preferences of each animal.

###### 1.2.2.1.2. Recipients’ individual training

Recipient animals were trained to poke to collect food rewards in the food magasin, by displaying a strong food-seeking behaviour (nose-poking repeatedly) (Supplementary Figure 3c-d). The individual training comprised different sessions with an increasing nose-poke ratio, starting from FR1 until FR5 nose-pokes in order to obtain a reward. Recipient animals were re-trained in this individual protocol during the social testing days (except for the first session), in order to avoid extinction of food-seeking behaviour, as during the PCT they were not in control of their own reward delivery anymore.

##### 1.2.2.2. Prosocial Choice Task protocol in the two-chamber arena

After individual training, mice were tested with the prosocial choice task in the two-chamber setup. Trials started when the focal nose-poked into the start-trial poke. The recipient could display food-seeking behaviour by nose-poking repeatedly into its nose-port, while the decision-maker could choose the top or bottom poke, which were counterbalanced between the pairs of animals to be prosocial and selfish choices. Choosing any of the options would always deliver a rewarding pellet for the decision-makers; however, the rewards for the recipient would only be available after a prosocial choice. Twelve animals were tested in the social task, in eleven – 40 minutes sessions. In four of these animals, data was only available for the first 4 sessions due to experimental problems. Mice underwent food-restriction 2h prior social testing.

#### 1.2.3. Hardware and peripherals

Both behavioural setups (double T-maze and double chamber) were custom-made built using laser cuter (Epilog Laser – 60W) and 3D printing (MakerBot Replicator 2 and Ultimaker 3), and progression of the task structure was controlled by pyControl (Akam et al., 2022). State-machine peripherals used from pyControl were adapted to fit our configurational needs. The latest modifications used are explained below.

##### Nose-ports (both setups)

Four pyControl nose-poke devices were used for controlling both decisions and food-seeking behaviour, two per individual maze. The IR-beam arms and LED were desoldered from the nose-poke boards and extended with wires so we could keep the components inside the setup and the boards outside the sound-attenuation box. These components were attached to a custom-made 3D printed nose-port piece which was attached inside the 1.2 cm inner walls that divided the different areas (Figure 1a-b)

##### IR detectors for detecting animal position to control behavioural state (maze setup)

Three pyControl nose-poke devices were used to detect the animal position inside the mazes to drive the behavioural state machine. In order to use them, the devices were modified. We desoldered the IR-beam arms, extended with wires to keep the components inside the mazes and the boards outside the sound-isolation box, and we also changed the IR emitter (TSUS5202, Vishay) to allow for a longer detection range given by the width of the corridors (6 cm for choice area and 7 cm for reward areas).

##### Sound attenuation box (both setups)

Each double T-maze was located inside a custom-made sound attenuation box. They were built on a 40x40 cm wooden cabinet with a single door. Sound isolation material (Regular Panel 60.2 Premiere, EliAcoustic) was placed in all inner walls, which provided 20 dBs sound attenuation. The inside of the isolation boxes was illuminated with dim white light (4 lux) and infra-red stripes located on the ceiling of the box. There was a 5x5cm ventilator placed on the middle of the back wall to regulate the inner temperature, and a 3cm diameter hole below it to pull all hardware wires out of the boxes. In this way we organised all wiring and electronic boards outside the boxes, to decrease possible audible and temperature interferences.

##### Custom-made pellet dispenser (both setups)

Food pellet rewards were delivered using custom-made pellet dispensers, which were built of a mix of 3D printed and laser cut parts, and actuated by stepper motors (NEMA 42HB34F08AB, e-ika electrónica y robótica, Spain) controlled by a stepper motor driver board. All of these were placed outside the sound attenuation box, to minimise the impact of the possible sound cues during the experiments. The palatable food pellets were dispensed to 3D printed food receptacles attached to the walls of the maze with magnets through a silicon tube that crossed the isolation box. Design files for the pellet dispenser and food receptacles can be found **here**.

##### Custom-made pneumatic doors (maze setup)

The sliding doors that control access to the different areas were made from 3mm transparent acrylic and built on top of a 3D printed piece containing a ball-bearing to allow smooth sliding of the door. These were actuated by pneumatic cylinders (Cilindro ISO 6432, Vestonn Pneumatic, Spain) placed below the base of the maze, providing silent and smooth horizontal movement of the doors. These were in turn controlled via solenoid valves (8112005201, Vestonn Pneumatic, Spain) interfaced with pyControl by using an optocoupled relay board (Cebek-T1, Fadisel, Spain), to prevent from possible electrical interferences coming from the solenoid valves coils. The speed of the opening/closing of the doors could be independently regulated by adjusting the pressure of the compressed air to the solenoid valves.

#### 1.2.4. Data acquisition systems and data analysis

##### 1.2.4.1. Video data

Individual training and experimental sessions were recorded with a high resolution infra-red sensitive camera (PointGrey Flea3-U3-13S2M CS, Canada) under infra-red illumination, capturing at 30fps with 1280x960 pixels resolution. Cameras were positioned centred above the setups to enable fine tracking of the animals’ position and pose estimation. Visual reactive programming software *Bonsai* (Lopes et al., 2015) was used to trigger the recording of the cameras and to start the behavioural task. It was also used to temporarily crop the experimental session videos into single trial videos during post-processing, using the IR light of the nose-ports that indicated the start of each trial and the choice moment.

##### 1.2.4.2. State machine and behavioural data

Behavioural data and general position of the animals in the automated mazes were extracted from pyControl software (Akam et al., 2022), and parsed with Python 3.0. For each session and animal, we extracted: number of trials, performance (trials/min), trial duration, prosocial preference (prosocial choices/total trials), frequency of nose-pokes, poke specificity, latency to decide. Moreover, for the experiments performed with the maze-like configuration, we extracted the latency to enter and to exit the reward areas after the automated doors opened.

##### 1.2.4.3. Pose estimation of mice and behavioural quantification

For tracking and pose estimation of the animals, we used the Bonsai-DeepLabCut interface (Python 3, DLC 2.2) (Kane et al., 2020). We first trained a ResNet-50 neural network for pose-estimation on labels of single animals taken from videos of a single individual maze (cropped image). Specifically, we labelled 426 frames from 5 videos/animals (95% was used for training with a 0.6 p-cutoff) for 600,000 iterations keeping the default network parameters. Then, a custom workflow of the Bonsai-DLC interface was used to batch-track the body pose of the interacting mice in all single trial videos from all PCT sessions. The workflow was able to simultaneously track the mice on any of the sides of the double maze, by applying a ROI for each side with an offset of the maze to maintain the original frame coordinates.

Finally, Python 3.0 scripts were written to analyse the social interactions happening during the experimental sessions. Tracking data from trials was split between prior and after choice period. We then extracted the coordinates position of body parts, which were used to compute different parameters to study the social dynamics during the social task. We extracted the location of the snouts of the mice and computed the euclidean distance between the animals and the distance in the X coordinate of the snouts from the central partition. Relative head-orientation of the two interacting animals was calculated by computing the angle-line from the label in between the ears of *animal A* and the label of the nose of *animal A*, minus the angle-line in between the ears of *animal A* to the nose label of the *animal B*. Furthermore, these variables were extracted in specific regions of interest such as the two nose-pokes in the choice area, the area around the wall dividing the two mazes in the choice area, and around both food-receptacles in the selfish and prosocial reward areas. Also, time spent in these ROIs was calculated.

### 1.3. Statistical analysis

Statistical analysis shown in this section will include the standard and widely used frequentist approach besides the Bayesian approach on the presented data, being the latter a convenient tool to discern those results showing evidence of absence of an effect from absence of evidence (Keysers et al., 2020).

Data extracted from the state-machine pyControl and the pose-estimation from DLC was parsed and processed with Python 3. We then used IBM SPSS Statistics 26 and JASP 0.16.2 to perform probabilistic and bayesian analysis on statistical differences between the extracted and studied variables.

<u>Prosocial preference</u>: repeated measures (RM) ANOVA with ‘session’ as within subjects factor was performed to assess the prosocial choice preference over the course of testing sessions (Figure 1d). In the 2-chambers setup, we further evaluated preference for the bottom nose-poke (Figure 8e), where proportion of choices towards the bottom choice was calculated over sessions using RM ANOVA the same way.

Prosocial Choice Index: we computed a prosocial choice index (PCI) to quantify individual differences on choice preference against chance over testing sessions,

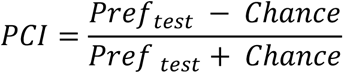

where *Pref_test_* corresponds to the proportion of prosocial choices during social testing sessions, and *Chance* is understood as the proportion of choices equal to 50%. The PCI values show the strength of change in prosocial preference from 50% preference for each mouse; [+] PCI show an increase on prosocial preference on social testing sessions compared to chance, [-] PCI show a decrease on prosocial preference from 50%, while the values close to 0 show no change. We performed a one-sample t-test to assess if the distribution of PCIs was different from chance (0) (Figure 1e, Figure 7b). For the case of the bottom preference (Figure 8f), we computed the PCI for the opposite preference of those mice that had the prosocial option on the higher poke.

<u>Permutation test to classify mice as prosocial, selfish or unbiased according to their</u> <u>preferences</u>: to address individual variability on prosocial preference, we performed a permutation test to identify those mice that showed significant change on choice preference against chance. For each animal separately, we generated a distribution of 10.000 permuted PCIs by shuffling the sequences of all choices during social testing with same-length sequences of choices with prosocial preference equal to 50%. Mice then were assigned to three different categories by comparing their actual PCI to the 95% confidence interval (CI) of the distribution of randomised indexes (mouse with actual PCI in 2,5% upper bound was considered as prosocial, mouse with PCI in 2,5% lower bound was considered selfish, and those mice with PCI falling inside the 95% were considered as unbiased). Lower and upper bound for each individual’s distribution of each experiment can be found in (Table 1).

<u>Side preference (individual training):</u> we used one sample t-test to check for differences in the side preference against chance (50) on the last two days of the individual training of mice for the sets of experiments. (Supplementary Figure 1c)

<u>Latency to decide:</u> we compared the latency, from trial onset to nose-poke (choice) at either the prosocial or selfish pokes, using paired samples t-test. (Figure 3b)

<u>Time exploring nose-pokes:</u> for focal and recipient animals we computed the time spent exploring the area around the prosocial and selfish nose-pokes per trial. For this, we computed the time that the DLC labels of the head and nose of both animals was detected in the two different ROIs around the two nose-ports, to ensure that we were measuring exploratory behaviour. This included the moments where animals were nose-poking, but also sniffing and investigating around the port, and was considered as a more global measure of investigation of the options. We used paired samples t-test to assess for differences between the conditions (i.e. prosocial or selfish ROIs). (Figure 2d and Figure 3c)

<u>Latency to visit reward areas:</u> we used paired samples t-test to assess differences in the averaged latency to enter into the prosocial and selfish reward areas after the choice moment, for focals and recipients. (Figure 2e and Figure 3d)

<u>Time exploring feeders:</u> we calculated the time that the head of focals and recipients was detected inside a ROI around the prosocial and selfish feeder (similar to that for the time exploring the nose-pokes) as a general measure for feeder investigation. We used paired samples t-test to assess for differences according to trial type. (Figure 2f and Figure 3e)

<u>Latency to leave reward areas:</u> we used related samples t-test to assess for differences in the latency to exit from the prosocial or from the selfish reward areas to start a new trial for both focals and recipients. (Figure 2g and Figure 3f)

<u>Pokes frequency:</u> we used paired samples t-test to assess differences between the frequency of pokes that recipients did to each type of nose-port per trial. (Figure 2b)

<u>Pokes specificity:</u> we calculated the specificity of the nose-pokes (n° prosocial pokes/ total n° pokes *100) done by recipients and used related samples t-test to assess differences according to trial type. (Figure 2c)

<u>Time in close distance</u>: using data from pose-estimation, we calculated the time that both animals spent in a distance less than 60 pixels (equivalent to nose-to-nose investigation). Paired samples t-test was used to evaluate differences according to trial type. (Figure 4b)

<u>Interaction time in partition:</u> we calculated the time that both animals spent together in a ROI around the partition in the choice area prior to decision. We used paired samples t-test to evaluate differences according to trial type. (Figure 4c)

<u>Social distance during interactions in partition:</u> we calculated the Euclidean distance between the nose label of focals and recipients during the interactions prior to choice in the partition ROI. Paired-samples t-test was used to assess for differences in this measure according to the trial type. (Figure 4d)

<u>Distance to wall during interactions in partition:</u> we calculated the distance of both focals and recipients towards the partition that separated both animals during the interactions prior to choice in the partition ROI. RM ANOVA was used to assess for differences in the distance to the partition of focals and recipients according to the trial type. (Figure 4e)

<u>Head orientation during partition interactions prior to choice:</u> we calculated the head orientation towards the other animal for both focals and recipients during the interaction in the partition of the choice area. We used independent samples t-test to assess for differences between them. RM ANOVA was used to assess for differences in the head orientation of focals and recipients according to the trial type (Figure 4f-g)

<u>Time interacting in feeders</u>: we checked the time that both animals spent interacting in the reward areas while being next to the feeders (ROI that comprised the feeders and the part of the corridor adjacent to the other animal) and used related samples t-test to check for differences according to the trial type. (Figure 5b)

<u>Head orientation during interaction in feeders</u>: we calculated the head orientation of both focals and recipients during the interaction time in each of the reward areas. We used independent samples t-test to check for differences between the angles of the animals according to their role. RM ANOVA was used to assess for differences in the head orientation of focals and recipients according to the trial type. (Figure 5c-d)

<u>Distance to wall during interaction in feeders:</u> we calculated the distance of both focals and recipients towards the partition that separated both animals during the interactions in the reward areas. We used independent samples t-test to check for differences between focals and recipients. RM ANOVA was used to assess for differences in the distance to the wall of focals and recipients according to the trial type. (Figure 5e-f)

<u>Individual differences between prosocial and selfish pairs:</u> we quantified (1) the total prosocial nose-pokes performed by recipient mice during the first session of PCT, (2) the latency of recipients from choice until entering into the reward zones, (3) the head orientation of focals during the displays of food-seeking behaviour of recipients, and (4) the distance to the wall of focals during the interactions in the selfish reward area. We then used one independent samples t-test to assess for differences in each of these variables comparing the extreme groups of the prosocial category (prosocial vs selfish animals). (Figure 6)

## Data availability

All data reported in this paper have been deposited at Zenodo and are publicly available as of the date of publication. DOI: 10.5281/zenodo.14628596

Any additional information required to reanalyse the data reported in this paper is available from the lead contact upon reasonable request.

## Author contributions

Conceptualization, C.M. and J.E.-A.; Methodology, C.M. and J.E.-A.; Hardware, J.E.- A. and V.J.R-M.; Formal Analysis, J.E.-A; Investigation, J.E.-A.; Resources, C.M.; Writing – original draft preparation, J.E.-A.; Writing– review & editing, J.E.-A and C.M.; Visualization, J.E.-A.; Supervision, C.M.; Funding acquisition, C.M.

## Acknowledgements

We thank the members of the Neural Circuits of Social Behavior laboratory for fruitful discussions and feedback on the results of the work.

---

**Supplementary Figure 1.**
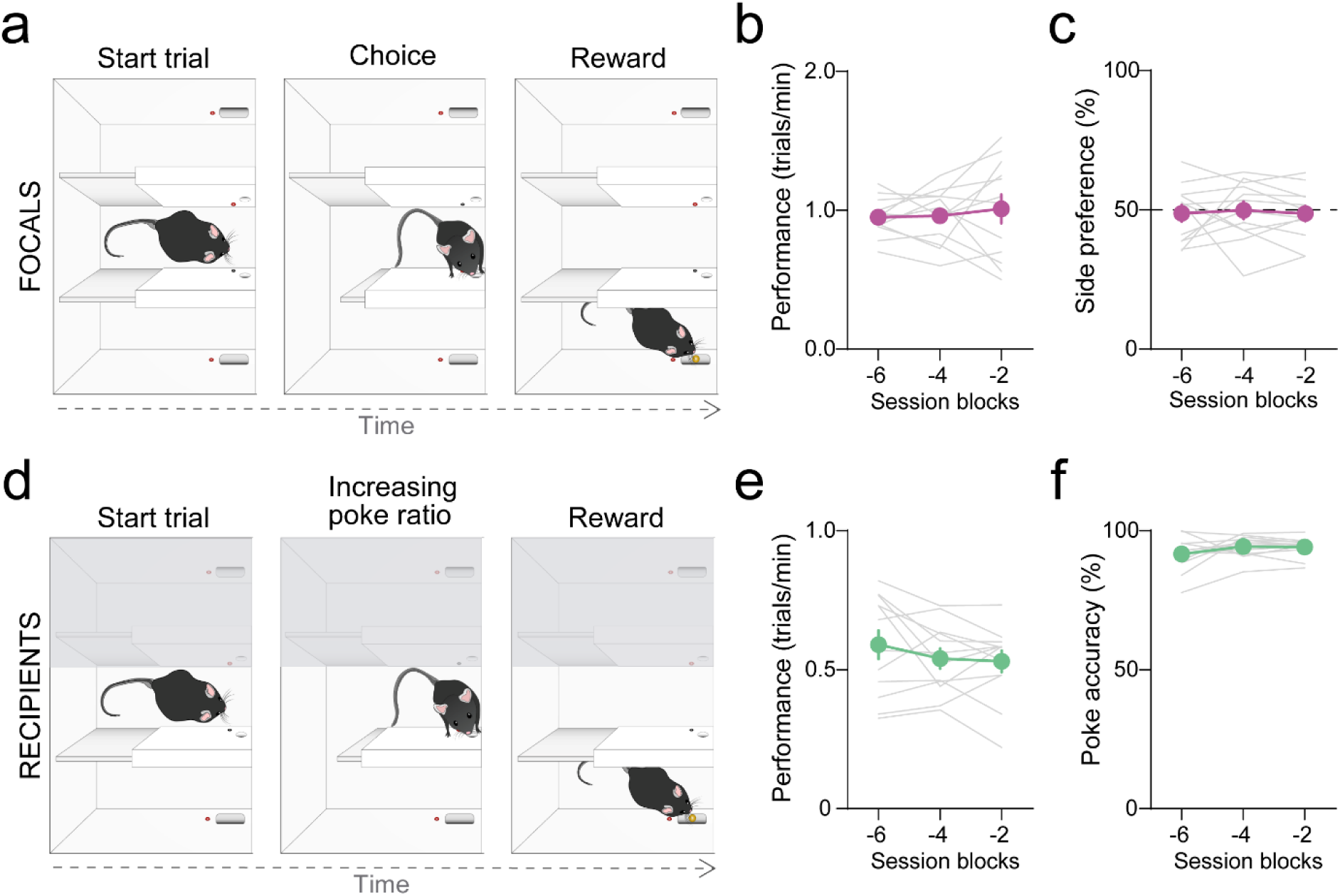
Individual training in the double T-maze before the PCT. **a**. Trial structure for focals’ individual training in the double T-maze, where mice choose between two pokes in the choice area to gain access to the corresponding reward area to obtain a pellet (both sides rewarded). **b**. Performance (number of trials divided by the session duration in minutes) of focal mice during last 6 sessions of individual training before the social task, averaged in blocks of 2 sessions. **c.** Side preference during last phases of individual training. Proportion of choices during last sessions of individual training to the side that will be prosocial in the PCT. Animals perform at chance. **d**. Trial structure for recipients’ individual training, where mice increase the poke ratio to gain access to reward only on one side of the maze. **e**. Performance of recipient mice during last sessions of individual training. Same as **b** for recipients. **f**. Nose-poke accuracy. Proportion of pokes towards the active nose-port over last sessions of individual training. Note that most of recipients poke almost exclusively into the port which leads to reward, which corresponds to the prosocial port during the social task.

**Supplementary Figure 2.**
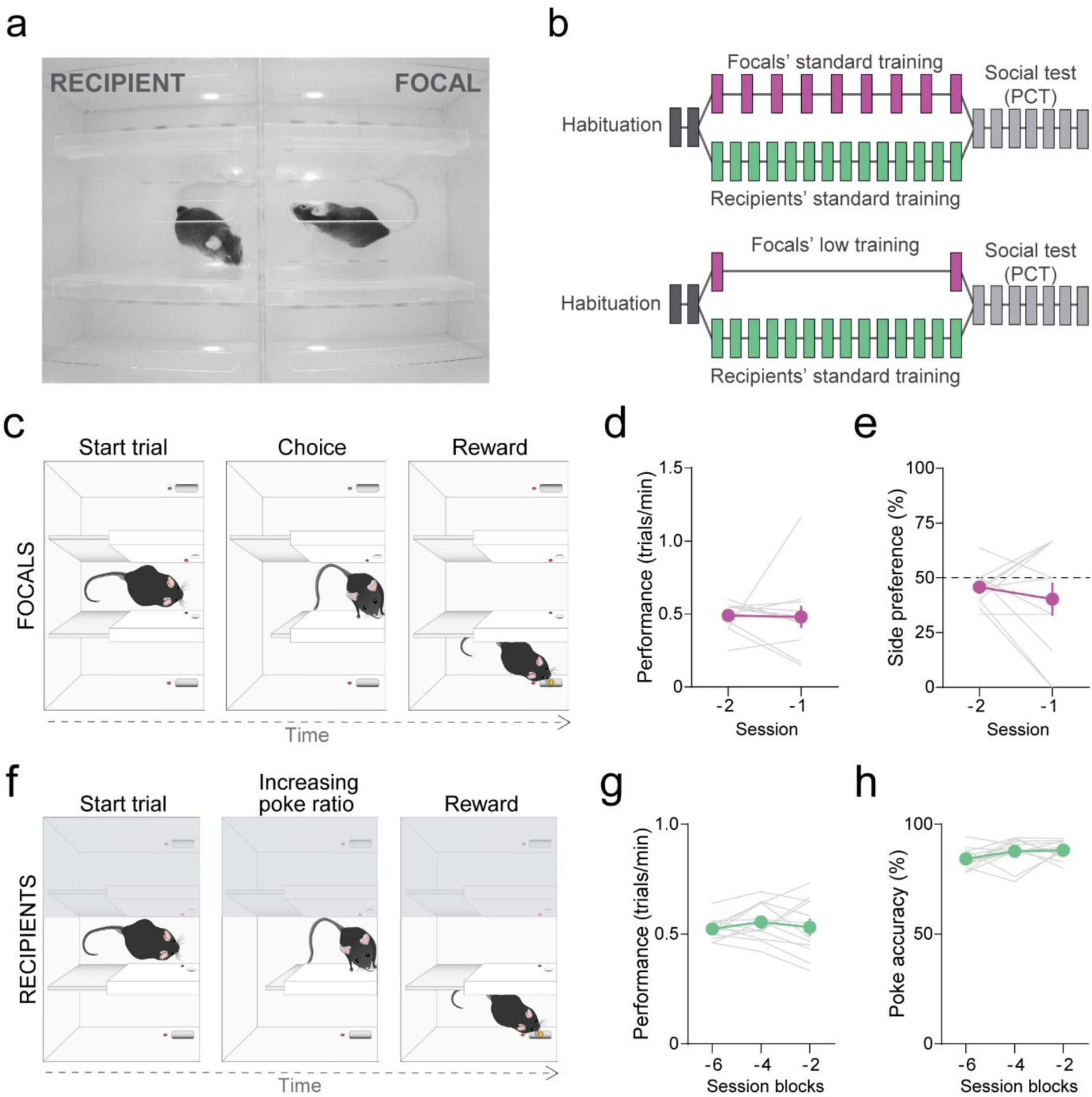
Low individual training protocol for focals in the double-T maze PCT. **a**. Real example image of mice in the choice area during a PCT session in the double T-maze setup configuration used for testing prosocial choices of focal mice with low-level training. **b**. Schema of standard individual training for recipients and low-level training for focals. Focals and recipients perform 2 sessions of habituation to the maze, then they all undergo a fixed-ratio 1 protocol session. Recipients continue their standard training protocol however, focals only perform an additional short fixed-ratio 1 session just prior to social testing. **c.** Focals training schedule, once in the choice area a single nose-poke into one side port will give access to reward in that side. **d.** Focals’ performance (number of trials/session time) for the only two sessions they were trained. Average of animals in purple. Individuals in grey. **e.** Side preference of the only two sessions focals were tested. Proportion of choices to the side that will be prosocial in the PCT. **f.** Recipients training schedule. **g.** Recipients’ performance (number of trials / session time) for the last six sessions before the PCT, averaged in blocks of two sessions. In green average of all animals, in grey individuals. **h.** Recipients’ nose-poke accuracy for the last six sessions before the PCT, averaged in blocks of two. Percentage of nose-pokes towards the prosocial side.

**Supplementary Figure 3.**
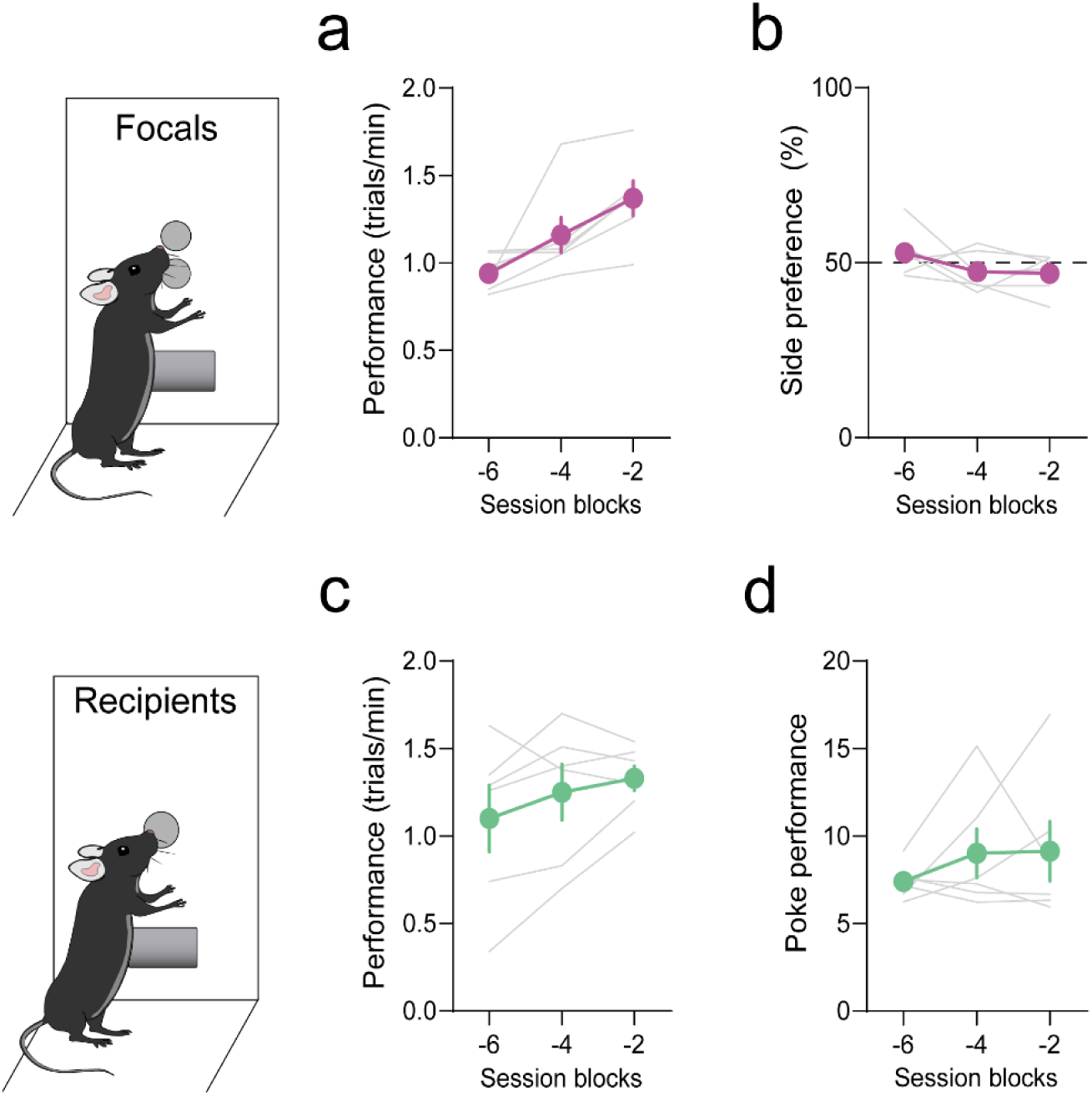
Individual training in the two-chamber before the PCT. **a**. Performance of focal mice during last phases of individual training, calculated as total trials/session duration in minutes. X-axis are the sessions before the PCT averaged in two (i.e., -2 is averaged data from the last two sessions of training before PCT). **b**. Side preference during the last phases of individual training. Proportion of choices during last sessions of individual training to the side that will be prosocial in the PCT. **c**. Performance of recipient mice during the last phases of individual training. Same as **a.** for recipient mice. **d**. Nose-poke performance during last sessions of individual training, calculated as total number of nose-pokes by number of trials. For all graphs: thicker line shows mean±SEM, thinner lines represent data from each individual.

**Supplementary Video 1. Example prosocial trial in the PCT with double T-maze structure**

**Supplementary Video 2. Example prosocial and selfish trials in the PCT with double chamber structure**

